# Predictive modeling of antibiotic eradication therapy success for new-onset *Pseudomonas aeruginosa* pulmonary infections in children with cystic fibrosis

**DOI:** 10.1101/2022.10.25.513740

**Authors:** Lucía Graña-Miraglia, Nadia Morales-Lizcano, Pauline W. Wang, David M. Hwang, Yvonne C. W. Yau, Valerie J. Waters, David S. Guttman

## Abstract

Chronic *Pseudomonas aeruginosa* (Pa) lung infections are the leading cause of mortality among cystic fibrosis (CF) patients; therefore, the eradication of new-onset Pa lung infections is an important therapeutic goal that can have long-term health benefits. The use of early antibiotic eradication therapy (AET) has been shown to eradicate the majority of new-onset Pa infections, and it is hoped that identifying the underlying basis for AET failure will further improve treatment outcomes. Here we generated random forest machine learning models to predict AET outcomes based on pathogen genomic data. We used a nested cross validation design, population structure control, and recursive feature selection to improve model performance and showed that incorporating population structure control was crucial for improving model interpretation and generalizability. Our best model, controlling for population structure and using only 30 recursively selected features, had an area under the curve of 0.87 for a holdout test dataset. The top-ranked features were generally associated with motility, adhesion, and biofilm formation.

**AUTHOR SUMMARY:** Cystic fibrosis (CF) patients are susceptible to lung infections by the opportunistic bacterial pathogen *Pseudomonas aeruginosa* (Pa) leading to increased morbidity and earlier mortality. Consequently, doctors use antibiotic eradication therapy (AET) to clear these new-onset Pa infections, which is successful in 60%-90% of cases. The hope is that by identifying the factors that lead to AET failure, we will improve treatment outcomes and improve the lives of CF patients. In this study, we attempted to predict AET success or failure based on the genomic sequences of the infecting Pa strains. We used machine learning models to determine the role of Pa genetics and to identify genes associated with AET failure. We found that our best model could predict treatment outcome with an accuracy of 0.87, and that genes associated with chronic infection (e.g., bacterial motility, biofilm formation, antimicrobial resistance) were also associated with AET failure.

## INTRODUCTION

Cystic fibrosis (CF) is the most common fatal genetic disease among individuals of European descent. This disorder is caused by a dysfunction of the cystic fibrosis trans-membrane conductance regulator (CFTR) gene, for which over 2000 mutations have been identified [1]. The loss of CFTR function can cause a suite of systemic, physiological problems, although the greatest impact is in the lungs where abnormal thick mucus accumulation in the airways inhibits bacterial mucociliary clearance and allows microbial pathogens to thrive [2,3].

Bacterial airways infections in CF patients typically occur early on in life and can be difficult to treat. At an early stage of the disease, *Haemophilus influenzae* and *Staphylococcus aureus* typically colonize the lungs, but later *Pseudomonas aeruginosa* (Pa) becomes highly prevalent, chronically infecting with up to 32% of adults with CF [4–8]. Chronic infection with Pa is associated with decline in lung function, leading to increased morbidity and mortality [9]. Improved clinical care and treatment have increased the quality of life and life expectancy for CF patients considerably in the last 50 years [10]. In Canada, adult CF patients now represent more than 62% of all patients [11], yet, bacterial infections, particularly with Pa, continue to pose a major threat. Consequently, the eradication of Pa infections early in life, delays the establishment of chronic infections and improves long term lung function [12], therefore, it is crucial to enhancing the quality of life of CF patients [13].

Antibiotic eradication treatment (AET) is the standard procedure for treating new onset Pa infections; although the protocol varies according to region and care facility, the success rate is relatively high, 20% to 40% of patients fail to clear the infection [14–16]. While there are likely many reasons for AET failure, the genetic makeup of the infecting Pa population is clearly an area of intense interest, and factors such as variation in the exopolysaccharide Psl have been shown to contribute to increased biofilm formation and tobramycin tolerance [17]. Despite this, there has not yet been a systematic study of the relationship between Pa genetic diversity and AET failure, or an attempt to predict the latter from the former.

Machine learning and related statistical genetic methods are now broadly used in microbiology to predict clinical outcomes, sources of infection, antibiotic resistance, and genetic variants underlying traits of interest [18–25]. While both supervised and unsupervised machine learning techniques have been useful in this context, there are several fundamental challenges to the application of machine learning in genomics. The first challenge is simply the scope of genomic diversity. Since each genetic variant (or input feature in the parlance of machine learning) is an independent variable, datasets inherently have very high dimensionality [26]. These features can be single nucleotide polymorphisms/variants (SNPs/SNVs), k-mers, unitigs (high-confidence, non-ambiguous contigs of assembled k-mers), or gene presence/absence. Dealing with high dimensionality data is non-trivial and if ignored will often lead to model overfitting. For this reason, it is often necessary to use feature selection or extraction techniques to avoid fitting random variation and non-informative features.

The second common problem in genomics is the lack of high-quality or high-confidence phenotypic data. This is particularly true for traits related to virulence, pathogenicity, and antibiotic susceptibility, since these traits are often measured in model or in vitro system that differ from the ‘natural’ target system both biotically (e.g., different host or microbial interactions) and abiotically (different environment and growth conditions).

Other challenges to the application of machine learning to genomics include predicting complex traits, whether these complexities are brought about through polygenic inheritance, epistasis, gene-by-environment interactions, variable penetrance, variable expressivity, genetic heterogeneity (i.e., genocopies), or phenocopies [27–30]. In addition, sampling biases, and non-independent evolutionary histories (i.e., population structure) among samples can result in hidden and complex covariation among input features [25,31–36]. Despite these challenges, machine learning and other statistical genomic approaches have proven to be extremely valuable tools for dissecting complex genomic data [23–25,37].

The aim of this work was to build a machine learning model to predict new-onset AET success or failure using whole genome sequence from Pa isolates cultured from new-onset infections in CF patients (prior to antibiotic treatment). We preformed Random Forest predictive modeling, which consists of an ensemble of decision trees, each providing an outcome based on different subsets of variable genomic unitigs. Random forest modeling has become increasingly popular in genomics, because it has shown high performance with small sample sizes, high-dimensional data, and complex data structures [38–40]. Additionally, random forest models allow input variables to be ordered according to their importance, which enabled us to identify which genomic variants have a strong influence on the outcome [41]. We used a nested cross validation (NCV) design [42], population structure control to control for non-independent evolution histories of the Pa strains, and feature selection to reduce the size of the input feature set and identify those variants that may be associated with AET failure [38,43].

## RESULTS

### Genomic and Phenotypic Diversity of Pa isolates

Bacterial isolates were retrieved from a cross-sectional study of CF pediatric patients (aged 0– 18 years) with new-onset Pa infections during the period 2011–2016 at the Hospital for Sick Children, Toronto, Canada [44]. A total of 440 Pa isolates were recovered from sputum collected from 70 patients prior to an inhaled/intravenous tobramycin multi-step protocol hereafter referred to antibiotic eradication therapy (AET) [15]. To assess within-patient diversity, one or more isolates were sequenced for each patient, depending on the number of colony morphologies recovered from the sputum sample. Sample isolates were labeled as eradication if the Pa infection was successfully cleared by AET or failure if AET was unsuccessful (Fig. 1A).

**Fig 1.**
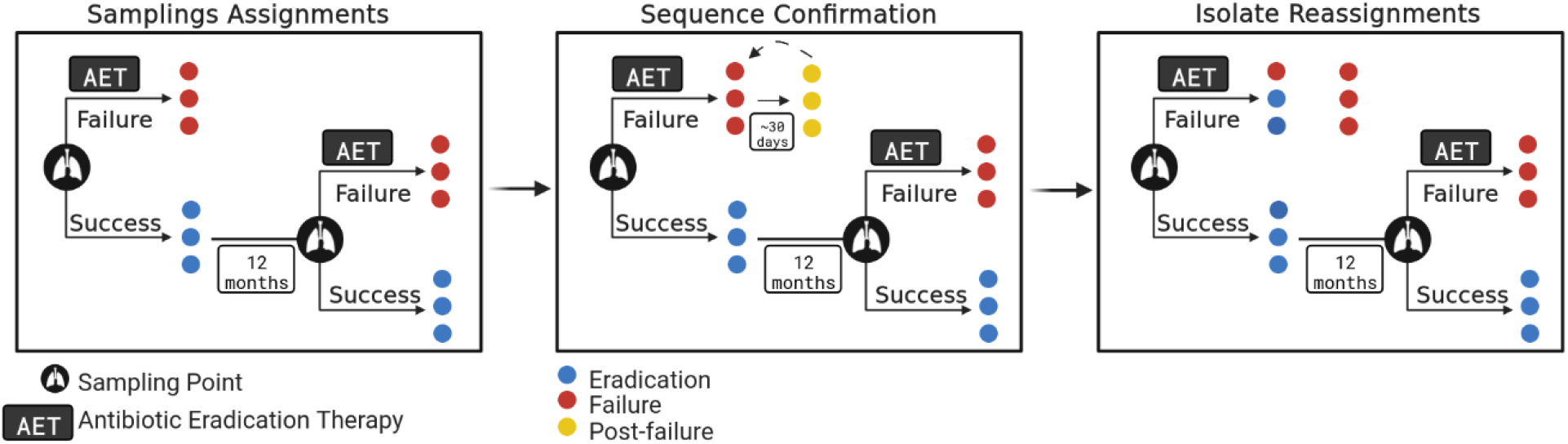
Overview of sampling strategy. (A) Sampling scheme for new-onset Pa infections and antibiotic eradication therapy (AET) with paths for treatment success or failure. New-onset Pa infections (identified by lung icon) were defined as first lifetime acquired infections or first infection 12 months after Pa clearance of the previous one. More than one isolate was collected if morphotype diversity was observed. AET (Inhaled/intravenous tobramycin multi-step protocol) either led to successful eradication (blue) or failure to eradicate (red). (B) Sequence confirmation and (C) sequence reassignment path for patients with AET failure. Persistent isolates (yellow) were collected, sequenced, and compared to pre-AET isolates in order to identify which clones survived AET. If a post-failure isolate was found to be nearly identical to a failure isolate, then that failure isolate would retain its failure label. If the post-failure isolate was not closely related to a prior failure isolate then those prior isolates were reassigned to the eradication success category.

New-onset infections usually refer to those acquired for the first time in the patient’s life. In this study, we also considered new-onset infections those acquired for the second or third time, but at least 12 months after the previous one. Among the 70 patients, five patients had a first infection instance that was cleared and a second one that failed AET, and 12 patients were infected more than once but AET was successful each time. In total, there were 90 infection episodes among the 70 patients, of these, 67 (74.4%) were eradicated and 23 (25.6%) failed. Of the 440 sequenced Pa isolates, 124 (27.2%) were recovered prior to AET failure and 316 (72.8%) were recovered prior to AET success.

We performed pangenome analysis to evaluate sample diversity within infections. While core genome distance between the samples was low (Fig. 2A), we found high levels of accessory genome variation among 15 eradication and seven failure infections episodes (Fig. 2B). Within-infection diversity could affect the outcome classification (i.e., eradication vs. failure) of individual isolates since the AET outcomes are assigned at the patient level. While this is not a problem when the infection was successfully cleared, it may be a concern in AET failure patients since the cause of treatment failure may be driven by only one of the multiple isolates recovered from that patient at that time. To confirm the AET outcome for each strain, we sequenced 55 isolates obtained after AET failure from ten patients (‘post-failure’ isolates, Fig. 1B), including six of the patients that showed within-episode accessory genome diversity (Fig. 2B, asterisk). We could not obtain post-failure isolates for two of the patients with high diversity infection episodes (SK006 and SK028). The post-failure isolates are identified with yellow labels in Fig. 1B and 2D.

**Fig 2.**
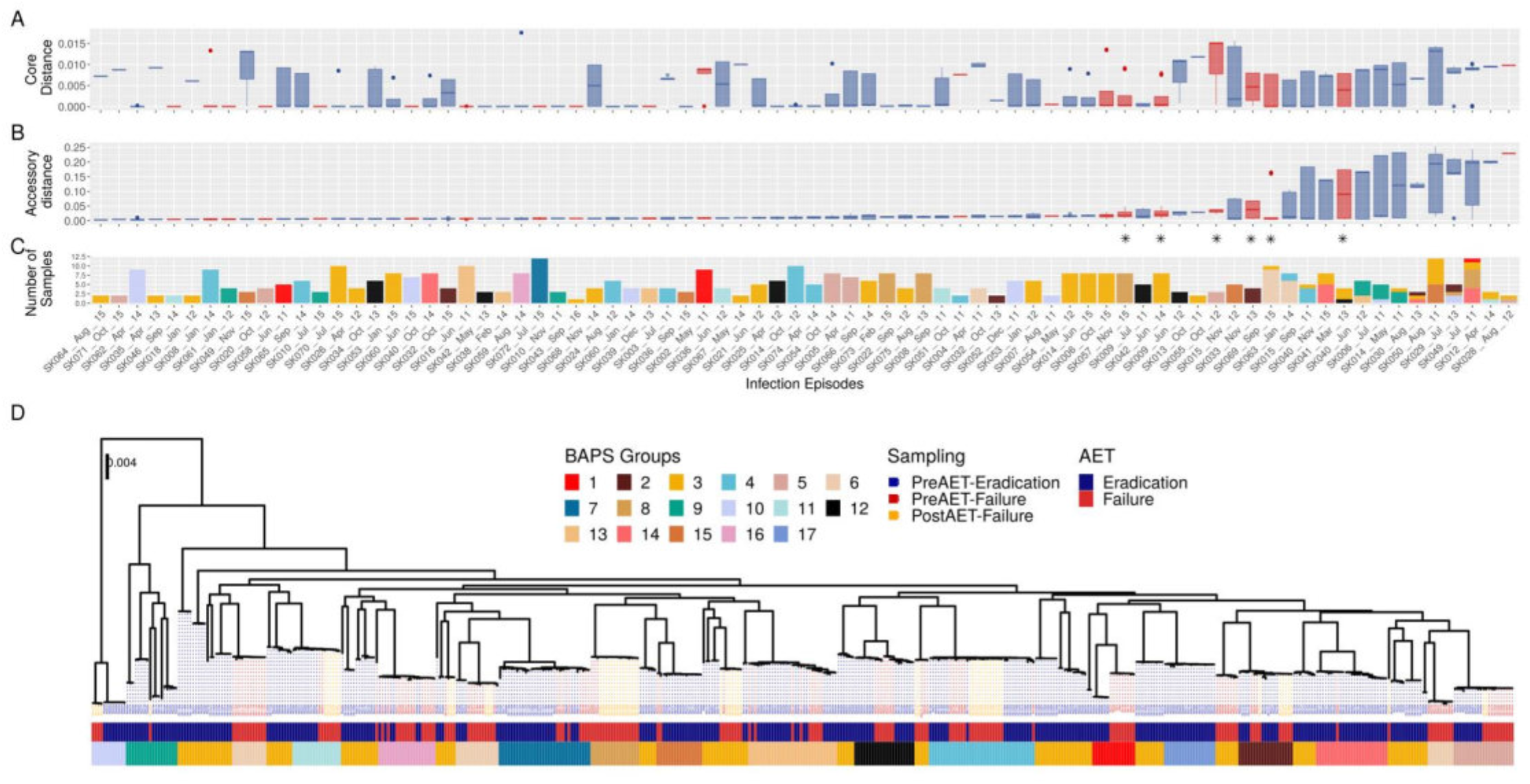
Genomic Diversity. (A) Core genome distance within each infection episode as estimated by MASH [100]. (B) Accessory genome distance (Bray-Curtis) within each infection episode estimated from ROARY analysis of gene presence/absence [97]. The infection episodes in both panels (A) and (B) are displayed in ascending order according to accessory genome distance median. (C) Number of isolates recovered in each infection episode and the proportion of those that correspond to each BAPS group. (D) Core genome, midpoint-rooted phylogeny of the evolutionary relationships between the 494 Pa strains. AET outcome is represented by the color coding of the dashed lines leading from the terminal nodes to the metadata rows, with blue showing AET success (i.e., eradication), red showing AET failure, and orange showing a post-failure isolate. The first annotation bar below the tree indicates AET outcome, while the second indicates the BAPS group.

We found that most post-failure genomes share a recent common ancestor with at least one pre-failure genome from the same patient based on their accessory genome distance (S1 Fig). We also evaluated the accessory genome distance between pre-failure and post-failure isolates, and ultimately, out of 28 pre-AET failure isolates, only three were reassigned from a failure to eradication phenotype based on accessory genome variability (S1 Fig). Given that there was very little genetic distance between non-reassigned pre-failure and post-failure genomes at both core and accessory genomes, and that sampling time after treatment was only approximately 30 days, we decided to include the post-failure isolates in the machine learning analysis (Fig. 1C). This had the benefit of increasing our sample size and ameliorated the difference between the number of eradicated and failed isolates (S1 Fig). The final sample set was therefore of 494 isolates, with 177 corresponding to AET failure samples (S1 Table).

Pa isolates in new-onset infections are usually acquired from the environment or upper-airway [45,46,47,48], therefore we expected high genetic diversity within the sample. To explore this, we built a core genome phylogeny including three reference genomes (PAO1, PA14, PA7) that are found in the three major Pa lineages previously identified by Ozer and colleagues [49]. We found that most of our isolates clustered in lineage 1, represented by the PAO1 reference genome, while a small number clustered in lineage 2, represented by the PA14 reference genome. None of our isolates clustered in lineage 3, represented by the PA7 reference genome (S1 Fig).

We observed that the AET success/failure outcome was highly correlated with the phylogenetic structure of the sample collection (Fig. 2D), with most of the clades containing isolates assigned to one of the two AET outcome phenotypes. This correlation between phenotypes of interest and the phylogenetic structure impacts machine learning predictions since it can introduce spurious associations between genomic diversity and the AET success/failure outcomes. This population structure bias, also known as population stratification or lineage effects, is caused by non-independent (i.e., correlated) evolutionary histories among strains in the sample [26]. Such biases have been extensively discussed in the context of both genome-wide association studies and machine learning predictive modeling [25,31–34,50–52].

We assessed and controlled for population structure using Bayesian Analysis of Population Structure (BAPS) [53–55]. The analysis supported the existence of 17 BAPS subpopulations or groups (Fig. 2C and D). Only two BAPS groups (14 and 17) were composed exclusively of AET success isolates, while the rest had representatives of both outcomes in different proportions. Most of the infection episodes were caused by homogeneous groups of isolates corresponding to a single BAPS subpopulation, with 76 of the 90 (84.4%) infection episodes caused by a single closely related clone (Fig. 2C and D). Counterintuitively, BAPS group 3 includes multiple distinct clades that span the entire tree. This classification reflects the genetic background of the strain collection and includes those clades that do not cluster into smaller distinct cluster, perhaps due to recombination (Dr. Gerry Tonkin-Hill personal communication) [56]. Of seven AET failures with high accessory genome variation, three were polyclonal, while of the 15 eradication episodes with high accessory genome variation, 11 were polyclonal (Fig. 2C). We also observed multiple closely related clades that included strains obtained from multiple patients, indicating that hospital transmission could be of importance [57,58].

Antimicrobial susceptibility testing (AST) was performed for 12 antibiotics for all the isolates via broth microdilution assays. We observed significant differences in the minimum inhibitory concentrations (MICs) between failed and eradicated isolates only for ciprofloxacin (CIP) and levofloxacin (LEV) (Fig 3). No association was found between tobramycin resistance and AET success/failure (Chi square, p value=0.09). Post-treatment isolates were also tested for antimicrobial susceptibility and found to have increased MIC levels for most of the 12 antibiotics tested (Fig. 3), revealing a rapid change in the antibiotic resistant ability of the Pa population during the infection. Only 13 isolates showed high levels of tobramycin resistance (MIC >256 μg/mL), with eight (61.5%) occurring in strains from AET failure samples and five (38.5%) from AET success sample. Resistance to tobramycin with MIC ≥ 16 μg/mL was found in 6.8% of the cases of AET failure and 1.6% of the AET success cases.

**Fig 3.**
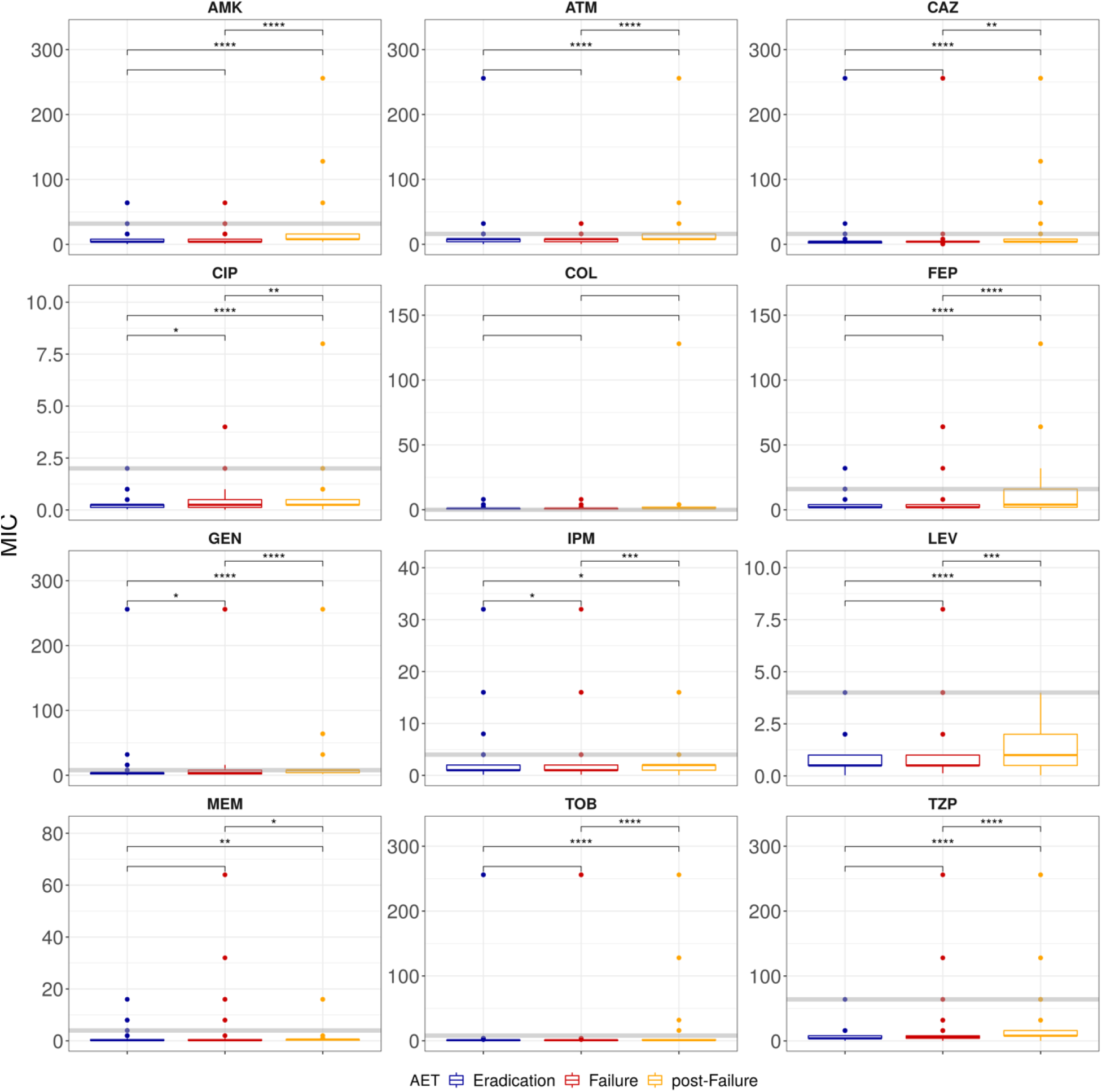
Antimicrobial susceptibility tests. Minimum inhibitory concentrations (MIC) levels of 12 antibiotics, for eradicated, failure and post-failure samples. The grey horizontal line indicates the breakpoint value. The acronyms correspond to AMK: Amikacin, ATM: Aztreonam, CAZ: Ceftazidime, CIP: Ciprofloxacin, COL: Colistin, FEP: Cefepime, GEN: Gentamicin, IPM: Imipenem, LEV: levofloxacin, MEM: meropenem, TOB: tobramycin, TZP: Piperacillin \+ tazobactam. Statical significance is represented with asterisks: ns: p>0.05,*: p<=0.05; **: p<=0.01;***: p<=0.001.

### Genome-Based Predictive Modeling of AET Success/Failure Outcomes

#### Input features

Input features representing the genetic variation of the sample population must be extracted from whole genome sequence data to build ML predictors. While most statistical genomic applications use SNPs, SNVs, or gene presence/absence data [19,21,59–61], short sequences such as k-mers or unitigs are becoming increasingly popular since they inherently incorporate polymorphisms, insertion/deletions, and gene presence/absence [62,63]. Here we used a presence/absence matrix of unitigs spanning the pangenome as input features to predict AET success/failure. Unitigs are high confidence contigs (i.e., assembled sequence reads) that have no mismatches or ambiguous residues. In addition to being of high confidence, unitigs also have the advantage of being less redundant than k-mers [62]. In order to reduce the number of input features, we selected a non-redundant set of unitigs by using only one representative unitig from set of identical patterns of presence/absence among the strain collection. We also removed unitigs with frequencies less than 5% and greater than 90% (reducing the number of features from 542,296 to 425,005) since these are highly unlikely to be strongly associated with our AET success/failure outcome, which was observed in 36% of the samples.

#### Machine learning predictor design

We used a random forest classifier with a nested cross validation (NCV) design to predict AET outcomes. NCV has a double loop analytical structure, with an inner loop for model/parameter selection, and an independent outer loop that assesses the quality. This approach maximized the use of our small number of samples, and enabled feature selection and hyperparameter tuning in a way that minimized the likelihood of model overfitting [42,64–66].

We employed population structure control (PSC) to control for correlated or non-independent evolutionary histories among our strains. PSC was implemented by clustering the strains by BAPS groups during the NCV outer loop. We then used BAPS group blocking, which assigns all strains in a BAPS group to either the training set or the test set, while maintaining class proportions (S2 Fig). We compared the performance of models with PSC and with no population structure control (nPSC) to evaluate the impact of lineage effects on AET outcome predictions.

We used recursive feature elimination (RFE) for model dependent feature selection [67]. RFE is a wrapper-type feature selection algorithm that ranks features by importance and recursively removes the least important ones until the selected number of features remain.

Finally, we used the combination of features selected in the different NCV iterations to fit a new random forest predictor. Essentially, the goal of this feature combination step was to find features that could generate accurate predictors independently of the train/test split; thereby, reducing biases in the model that can be particularly challenging when the number of samples is small, and the outcome phenotype is highly correlated with the phylogeny.

Given the imbalanced nature of the data we chose the area under the ROC curve (AUC) as the performance measure. Train and test AUC can be compared to evaluate over-fitting and population structure impact. We also used precision, recall, and the F1 score for the positive (AET failure) and negative (AET eradication) classes to assess model performance. Having a high recall of the positive class is extremely important because it reflects a low number of false negatives or a small type II error, which we are particularly interested in reducing (reduce the number of false eradication predictions) (S3 Fig).

Models built using an excess of features relative to the number of samples can suffer from overfitting and achieve poor generalization [25,43]. Consequently, we elected to try two approaches to reduce the number of features to improve model performance, complexity, and interpretation while reducing computational cost: 1) feature transformation via multivariate dimensionality reduction, and 2) feature selection involving filter and wrapped methods. The overall strategy was similar in both approaches and discussed in more detail below (Fig. 4).

**Fig 4.**
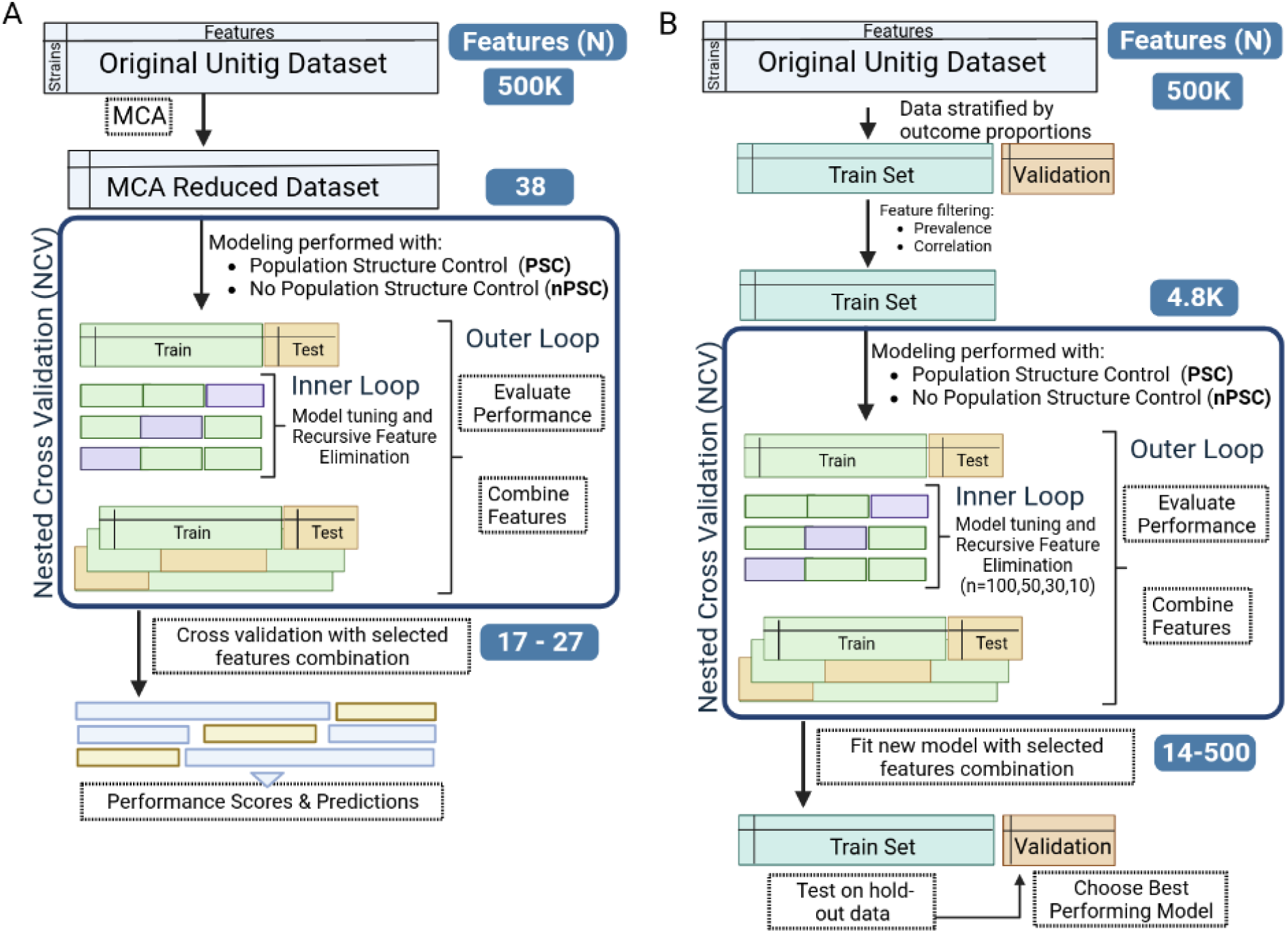
Machine learning overview. (A) Workflow for the prediction of AET outcome using MCA feature transformation. The original dataset (all one-hot-encoded unitig features found in a strain and the AET outcome of success or failure for that strain) was reduced via MCA to 38 dimensions used for downstream analysis. Nested cross validation (NCV) was used for model tuning and recursive feature elimination (RFE) to further reduce the number of features. Separate modeling was done with no populations structure control (nPSC) and population structure control (PSC), which was implemented by blocking the data based on BAPS groups. All the selected features from the NCV outer loop were combined for final model construction. The performance of the final models was evaluated via cross validation. (B) Workflow for the prediction of AET outcome based on unitig diversity. The original one-hot-encoded unitig data was divided into a training and a validation set stratified by the outcome proportions. Feature correlation filters were applied to remove redundant features, thereby reducing the feature dimension from >500k to 4800. Then we used NCV to further reduce the number of features in a model dependent manner with RFE in the same manner as discussed above. The feature combinations were used to fit a new predictor on the training set and the model evaluated on the hold-out validation set.

#### Machine learning predictor using multiple correspondence analysis (MCA)

MCA [68] was able to capture 80% of the variation in our unitig dataset in 38 dimensions, with the first two dimensions represented 12.3% of the total inertia (variation). While dimensions 1 and 2 did not clearly separate AET outcomes (Fig. 5A), they did more clearly distinguish BAPS groups (Fig. 5B).

**Fig 5.**
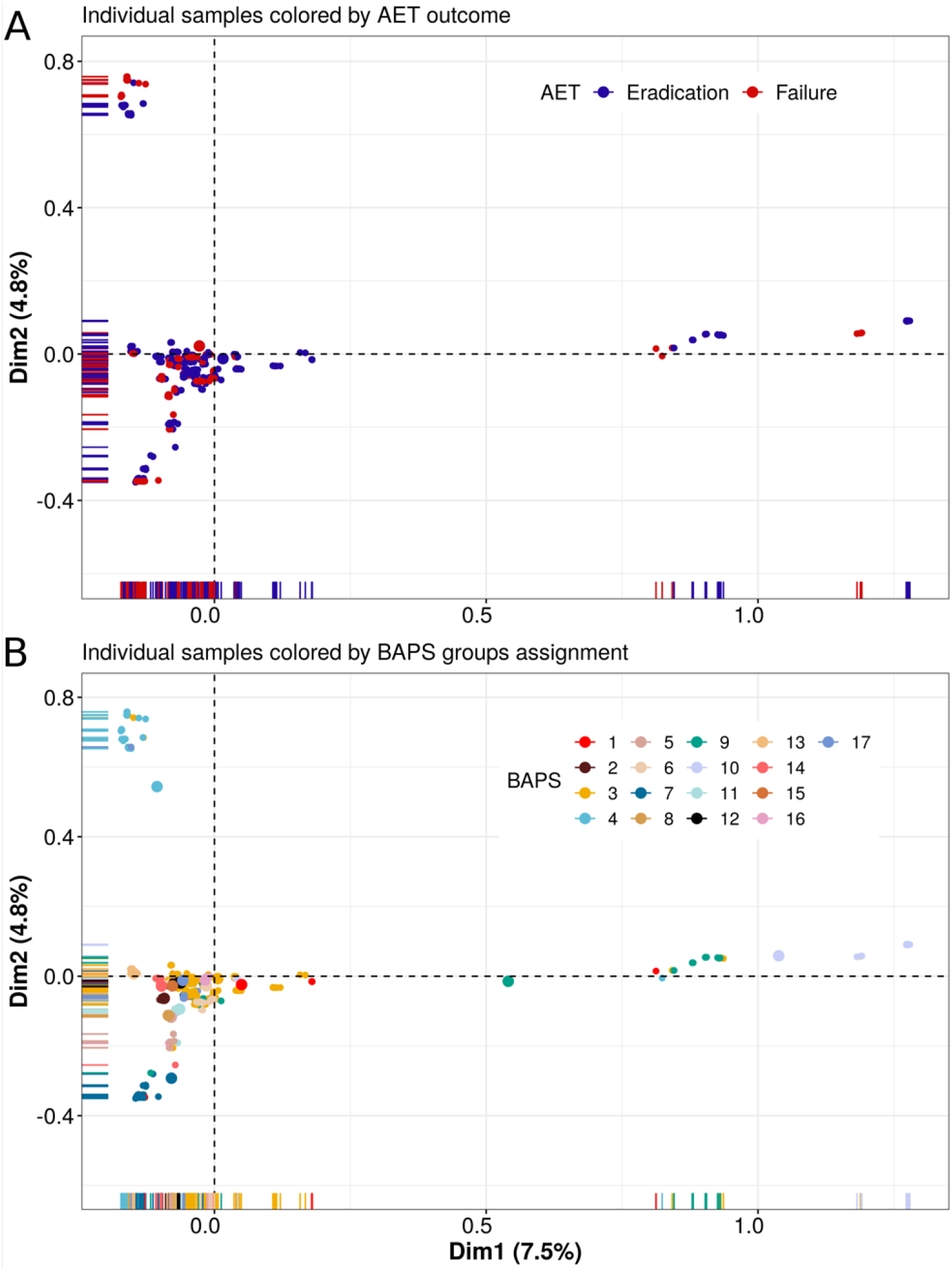
Multiple corresponding analysis. (A) The first two MCA dimensions (with percent variation explained) showing the clustering of 494 samples colored according to treatment outcome. (B) The same figure but with samples colored according to the BAPS group. Much of the variation captured in both axes is due to population structure of the sample.

The performance of the MCA-based random forest classifier was highly sensitive to PSC as demonstrated by a large difference in AUC values between train and test datasets (mean train AUC=0.96, mean test AUC=0.52, Fig. 6A, left panel). Low performance during testing indicate that the predictor failed to generalize to BAPS groups that were not included in the training set (Fig. 6A, left panel). On the other hand, when PSC was not applied, only a slight difference between train and test AUC was observed (mean train AUC= 0.98, mean test AUC= 0.88), the latter, however, shows higher variation (train AUC sd = 0.03, test AUC sd = 0.1), which is characteristic of small data sets, where different train/test splits impact the performance of the model (Fig. 6A, right panel).

**Fig 6.**
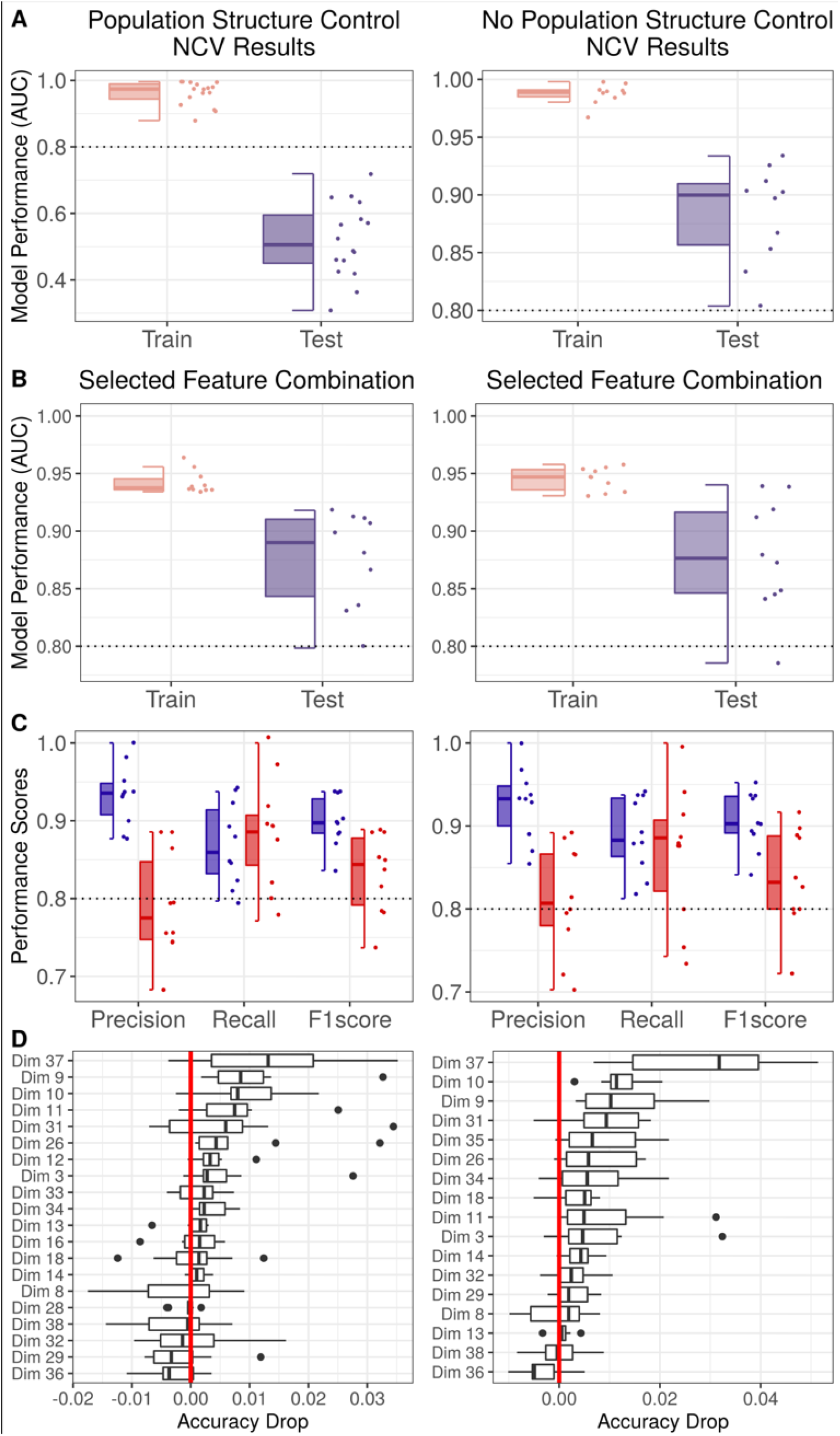
Model performance using MCA feature reduction. (A) Area under the curve (AUC) for training and test sets obtained during the nested cross validation (NCV) analysis. The left panel shows results with population structure control (PSC), while the right panel shows with no population structure control (nPSC). The same set of input features were used for both models (38 dimensions representing 80% of the variation). The impact of including PSC significantly lowers AUC values during testing. Note the different scale of the axes for the PSC and nPSC results. The dashed line at 0.8 provides a common reference throughout. (B) Train and test AUC obtained with cross validation using the reduced set of features previously selected. Performance is similar with and without PSC (27 features selected with PSC and 17 without PSC). (C) Precision, recall, and F1 score for the AET outcomes (blue indicating AET success, i.e., eradication, and red indicating AET failure) obtained via cross validation using a reduced set of features previously selected. (D) Boxplots showing the feature importance measured as accuracy drop in the model during the feature permutation importance analysis.

We performed feature reduction by selecting the ten most important features from each NCV outer split, resulting in 17 dimensions for the nPSC pipeline and 27 for the the PSC pipeline. Sixteen of the features selected in nPSC were also selected when PSC was applied. Both reduced set of features were used to train new predictive models so that we could compare the performance of features combinations obtained with and without PSC. The model trained with features selected from PSC performed as well as the one trained with the selected features from nPSC (Fig. 6B), as expected given that the difference between both sets of input features was minimal. Precision and recall scores were also similar for the two models (PSC: Mean Recall, Failure=0.86 and Eradication=0.88, Mean Precision, Failure=0.79 and Eradication=0.93. nPSC: Mean Recall, Failure=0.86 and Eradication=0.88, Mean Precision, Failure=0.81 and Eradication=0.93). Mean recall for the positive class (AET failure) was slightly better for the nPSC selected features model (Fig. 6C). The performance scores are summarized in Tables 1 and 2 for each step. The ranking of features by importance was very similar for both models and the features selected were robust as shown by the results of a permutation test (Fig. 6D). Dimensions 37, 10, 9 and 31 stand out as the most important in both approaches. The two pipelines used provide a similar final result despite the inclusion of PSC.

**Table 1:** MCA Model Performance

| Model <sup>1</sup> | Model Stage <sup>2</sup> | N Features | Train AUC (sd) <sup>3</sup> | Test AUC (sd) |
| --- | --- | --- | --- | --- |
| PSC | NCV | 38 | 0.96 (0.03) | 0.52 (0.10) |
| PSC | CV | 27 | 0.94 (0.01) | 0.88 (0.04) |
| nPSC | NCV | 38 | 0.99 (0.01) | 0.88 (0.04) |
| nPSC | CV | 17 | 0.94 (0.01) | 0.88 (0.05) |

<sup>1</sup> PSC, population structure control; nPSC, no population structure control
<sup>2</sup> CV, cross validation; NCV, nested cross validation
<sup>3</sup> AUC, area under the curve

**Table 2:** MCA Classification Report

|  |  | AET Success | AET Failure |
| --- | --- | --- | --- |
|  | N Samples | 64 | 35 |
| PSC | Precision | 0.93 (0.03) | 0.79 (0.06) |
|  | Recall | 0.87 (0.05) | 0.88 (0.07) |
|  | F1 score | 0.9 (0.03) | 0.83 (0.05) |
| nPSC | Precision | 0.93 (0.04) | 0.81 (0.07) |
|  | Recall | 0.89 (0.05) | 0.87 (0.08) |
|  | F1 score | 0.91 (0.03) | 0.84 (0.06) |

While MCA drastically reduced the number of features, yielding uncorrelated features, and generally provided good performance, this approach lacked interpretability and reproducibility, since the dimensions do not have any real meaning, but instead are linear combinations of the input variables. Consequently, we cannot easily use these features to identify variants associated with AET outcomes and explore causality. Additionally, analyzing new samples with these models can be quite complex.

#### Machine learning predictor using unitigs as input features

As an alternative to MCA-based feature selection, we directly used one-hot encoded unitig variation as input for the random forest predictor and performed feature reduction through a combination of filters and wrappers. Unlike our previous approach, in this case we divided our data into a training and a hold-out validation set (90% / 10% split respectively), maintaining class proportions, before performing the NCV step (Fig. 4B). We used a chi-squared association test to remove features with no association with the AET outcome and identified and removed highly correlated features (`Pearson correlation coefficient` > 0.7) in the training data set. This reduced the number of features from 425,005 to 4800.

We ran four independent pipelines selecting for different numbers of features during the RFE step (n features = 10, 30, 50, and 100) to account for the random nature of the algorithm and the impact of multiple feature combinations on the performance of our predictor [67]. The features selected in each NCV iteration outer loop were combined to fit a new predictor on the original training set and the performance tested on the hold-out validation set (Fig. 4B).

The effect of including PSC was substantial (Fig. 7A-left panel). Model performance varied widely between iterations. Notably for some of the splits we obtained very poor training performance (Mean Train AUC = 0.75, sd = 0.16). Furthermore, testing performance was very low (AUC < 0.7), which means that even when the learning process showed high performance, the generalization power was low (Mean test AUC=0.51, SD test AUC=0.03). This behavior is maintained despite the number of features selected in independent RFE runs. In contrast, when PSC is not applied, model performance is consistently high across splits (Mean train AUC = 0.97,sd = 0.03), both for training and testing (Mean test AUC = 0.91, sd = 0.04). Still, train/test AUC differences indicated that the model was overfitting when the number of features selected during RFE was set higher than 10 or 30 (Fig. 7A-right panel).

**Fig 7.**
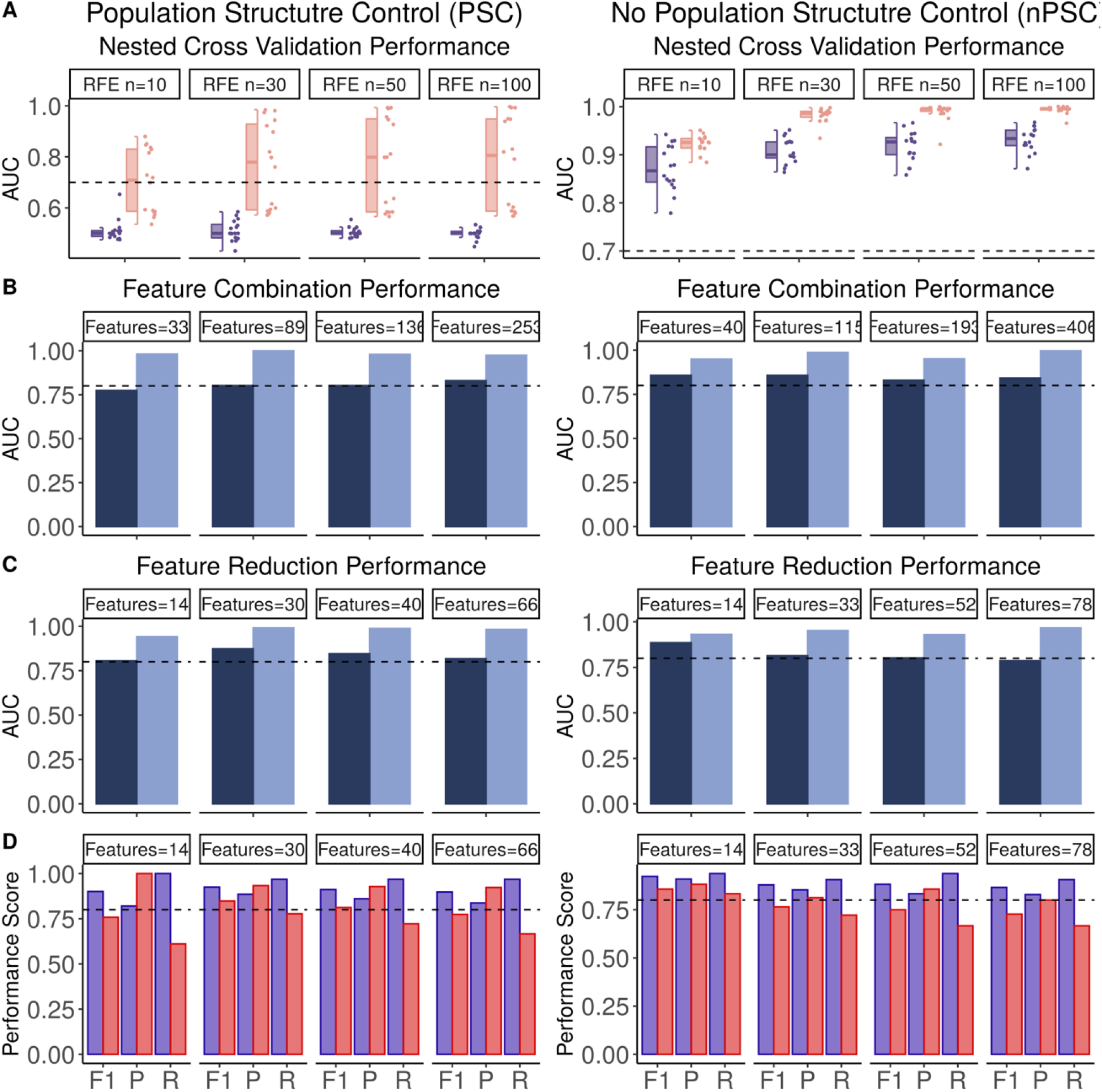
Model performance using recursive feature elimination. (A) AUC values for the training and test dataset obtained during the NCV analysis with different numbers of features selected (number of selected features = 10, 30, 50, and 100) during the recursive feature elimination (RFE) step. Left panel shows results with PSC, while right panel shows results with nPSC. Note that the y-axes are on difference scales. The dashed line at 0.7 indicates the threshold for feature selection. (B) Train and validation AUC obtained using the reduced set of features previously selected. (C) Train and validation AUC obtained using a further feature size reduction based on the training feature importance. (D) Precision (P), recall (R) and F1 score (F1) for the AET outcomes (blue indicating AET success, i.e., eradication, and red indicating AET failure) obtained using a further feature size reduction based on the training feature importance. The dashed line at 0.8 provides a common reference throughout B, C, and D.

We combined all of the features selected during the NCV iterations to train a new predictor using the entire training set, and then did performance testing using the hold-out data (Fig. 4B). For the PSC pipelines, features were only extracted from iterations with train AUC values above 0.7. Models trained with the feature combinations achieved high performance irrespective of the whether PSC was used during the selection process (Train AUC > 0.8, Validation AUC > 0.75) (Fig. 7B). However, there were signals of overfitting, particularly when using large feature combination sets. Tables 3 and 4 provide detailed performance scores for each step.

**Table 3:**
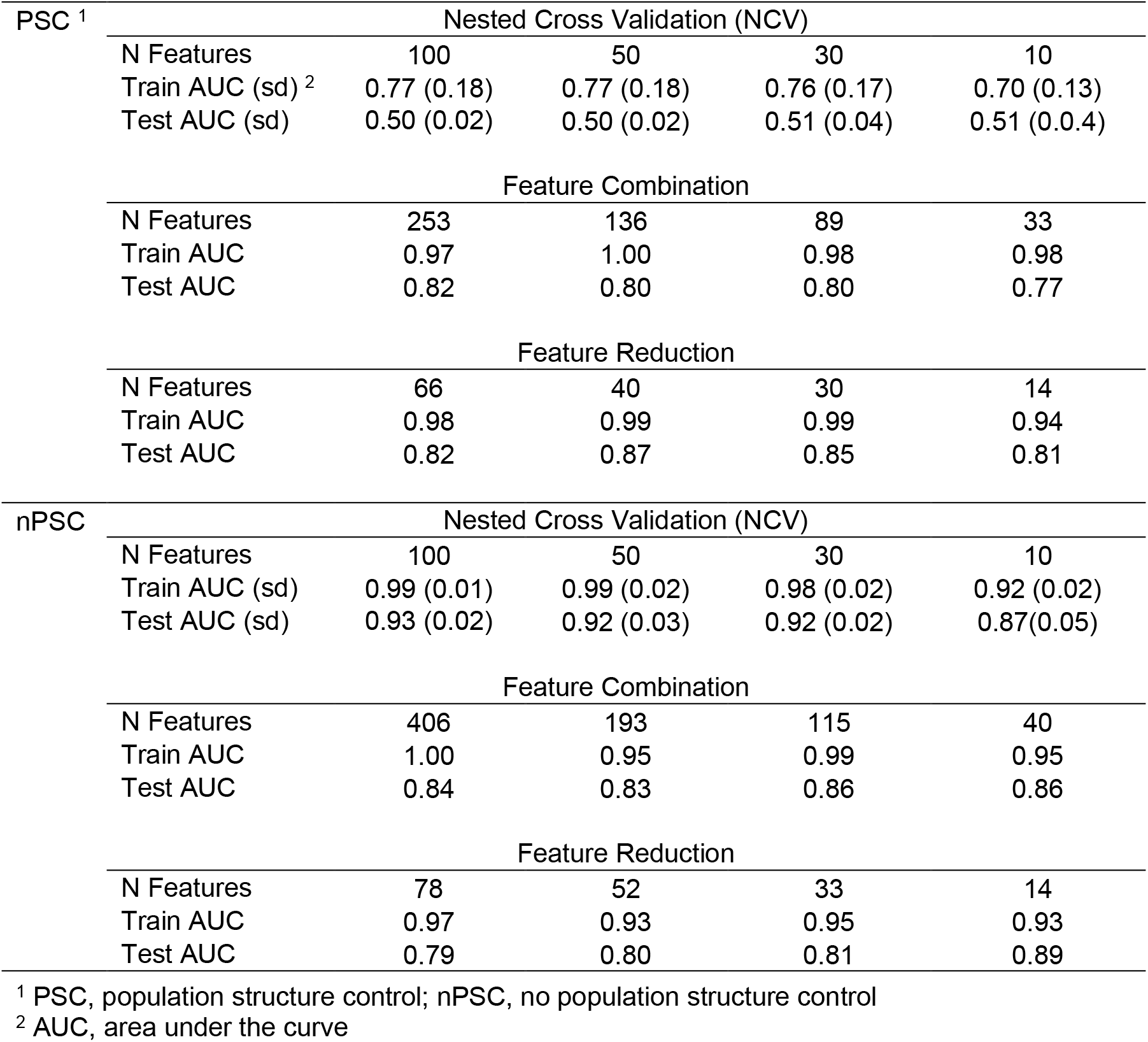
Unitig Model Performance

**Table 4:** Unitig Classification Report

| PSC <sup>1</sup> | N Features | 66 |  | 40 |  | 30 |  | 14 |  |
| --- | --- | --- | --- | --- | --- | --- | --- | --- | --- |
|  | AET <sup>2</sup> | Success | Failure | Success | Failure | Success | Failure | Success | Failure |
|  | Precision | 0.84 | 0.92 | 0.86 | 0.93 | 0.88 | 0.93 | 0.82 | 1.00 |
|  | Recall | 0.97 | 0.67 | 0.86 | 0.93 | 0.89 | 0.93 | 1.00 | 0.61 |
|  | F1 Score | 0.90 | 0.77 | 0.91 | 0.81 | 0.93 | 0.85 | 0.90 | 0.76 |

| nPSC | N Features | 78 |  | 52 |  | 33 |  | 14 |  |
| --- | --- | --- | --- | --- | --- | --- | --- | --- | --- |
|  | AET | Success | Failure | Success | Failure | Success | Failure | Success | Failure |
|  | Precision | 0.83 | 0.80 | 0.83 | 0.86 | 0.86 | 0.81 | 0.90 | 0.88 |
|  | Recall | 0.90 | 0.67 | 0.94 | 0.67 | 0.91 | 0.72 | 0.94 | 0.83 |
|  | F1 Score | 0.87 | 0.72 | 0.88 | 0.75 | 0.88 | 0.76 | 0.92 | 0.86 |

<sup>1</sup> PSC, population structure control; nPSC, no population structure control
<sup>2</sup> AET, antibiotic eradication therapy

#### Feature selection using feature importance

We eliminated features not associated with AET outcome based on their Gini Importance calculated during training. Gini Importance, also known as Mean Decrease Impurity, is a measure of how well a feature splits samples with different outcome or create groups with similar responses within. The reduced feature set was then used in subsequent rounds of predictor training. In some cases, the removal of feature did not have a major impact on performance, while in other cases it helped reduce the difference between train and test AUC (Fig. 7C). Based on train/validation performance differences and the recall and precision scores for both classes, the best model performance was achieved with 30 features obtained with the PSC pipeline (Train AUC = 0.99, Test AUC = 0.87, Precision: failure=0.93, eradication=0.88, Recall: failure=0.78, eradication=0.96), and a combination of 14 features obtained with nPSC (Train AUC = 0.93, Test AUC = 0.88, Precision: failure=0.88, eradication=0.91, Recall: failure=0.83, eradication=0.94) (Fig. 7D).

Although similar performance can be achieved with and without the inclusion of PSC, the features selected with the two approaches differed (S4 Fig). A significant number of features were selected consistently across the independent RFE instances within each pipeline, although there was very little overlap when comparing features from PSC and nPSC. For example, we found that only one feature was shared between the best performing models.

#### Testing the predictive power of randomly selected features

Given that similar performance can be achieved with different sets of features, we compared the performance of our best model to 500 models trained on 25 random features selected from the 4800 non-correlated input variables (S5 Fig). We found that most random groups of features lead to lower performance than our pipelines, although some combinations provide very good performance that even outperformed our more rigorous selection procedure discussed above. This surprising result is likely due to many of features being strongly correlated with the phylogeny; thereby, supporting the use of PSC to reduce lineage effects.

#### Assessing the impact of population structure control (PSC)

To determine if PSC could reduce lineage effects, we mapped the correctly and incorrectly predicted samples from the best-performing models onto the phylogeny (Fig. 8A). Phylogenetic mapping showed that the nPSC pipeline errors were primarily concentrated in clades with closely related strains that had different AET outcomes, whereas PSC is able to correctly predict these cases. This is an important finding since it indicates that features selected with PSC contain information that is independent of the phylogeny and thus should provide higher generalization power. Nevertheless, we cannot assign any causal relationship between the features we selected and the AET outcome unless we can distinguish between features that predict kinship and features that predict AET independently of phylogeny.

**Fig 8.**
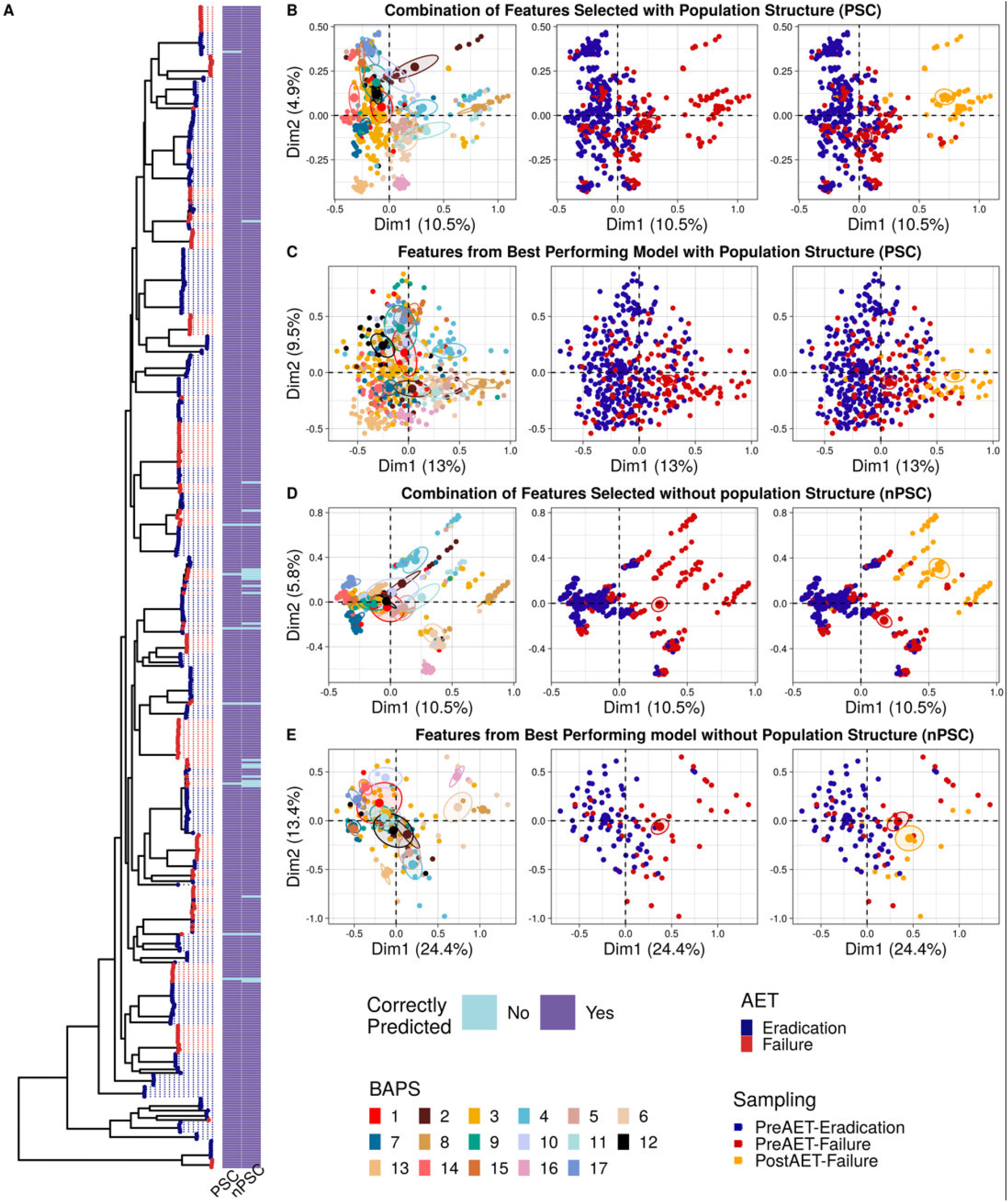
Assessing the impact of population structure control. A) Core genome phylogeny with tips colored according to AET outcome. The first annotation bar corresponds to predictions obtained with the best performing model of the PSC pipeline (purple = correct prediction, blue = incorrect prediction). The second annotation bar correspond to predictions obtained with the best performing model of the nPSC pipeline. (B-E) Multiple Correspondence Analysis (MCA) plots showing the clustering of the 494 samples based on the presence/absence pattern of features (i.e., unitigs). The three MCA plots in each row shows samples colored by BAPS group, AET outcome, and sampling category respectively. (B) PSC with all features selected during recursive feature elimination (RFE). (C) PSC with the 30 best performing features. (D) nPSC with all features selected during RFE. (E) nPSC with the 30 best performing features.

We used MCA to evaluate how well the feature combination selected by the PSC and nPSC pipelines can delineate AET success and failure. First, we analyzed the combination of features selected with the different RFE instances within each pipeline (a total of 253 unique features were selected with independent RFE for PSC and 406 for nPSC, Fig. 8B and D). We found that the first dimension can separate the phenotypes, but most of the variation is due to the persistent samples. This likely indicates that the persistent (post-failure) isolates carried unique genetic signatures that increased in frequency after treatment. The BAPS groups could also be clearly identified using both sets of features irrespective of PSC inclusion.

We then performed the same analysis but using the features from the best-performing models, that is, 30 features selected with PSC and 14 features with nPSC (Fig. 8 C and E). In both cases the first dimension could split both phenotypes very clearly, but the BAPS groups were less strongly delineated when using the PSC combination. This indicates that the PSC pipeline did a better job in creating a feature dataset that was population structure independent, and therefore, better suited for finding causal variants underlying AET failure irrespective of the clonal background. We also found that this reduced feature set diminished the impact of the persistent samples.

Individual unitig patterns showed low correlation values with the AET outcome both in the PSC and nPSC groups of features (max absolute values 0.36 and 0.4 respectively) (S6 Fig). This likely indicates that there are multiple evolutionary routes to persistence in the lung during early stages of the infection. Furthermore, some of those features were clearly associated to persistent samples, with most of the isolates in that group sharing the same pattern. This is likely due to differential fitness and survival of certain variants after treatment failure, which is consistent with our earlier finding that persistent isolates had increased antibiotic tolerance compared to the isolates obtained prior to AET.

### Genes Associated with AET Success/Failure Outcomes

Our best performing PSC model used 30 input features, while the nPSC model used 14 features (Fig. 9A and B). While we initially selected only one representative unitig out of a covarying set of identically or nearly identically distributed unitigs, we used all unitigs in the covarying sets for the functional annotation. When we included all identical and highly correlated unitigs, our best performing sets increased to 190 and 125 features for PSC and nPSC models respectively. We then mapped these expanded unitig sets to the Pa genome and recorded the overlapping coding sequence. When unitigs mapped to non-coding regions, we recorded the genes located both upstream and downstream from the mapping region. The number of features in coding and non-coding regions showed no significant difference between PSC and nPSC (Chi-squared test, p-value=1.0). The final list of genes associated to AET success/failure was of 128 for PSC and 84 for nPSC.

**Fig 9.**
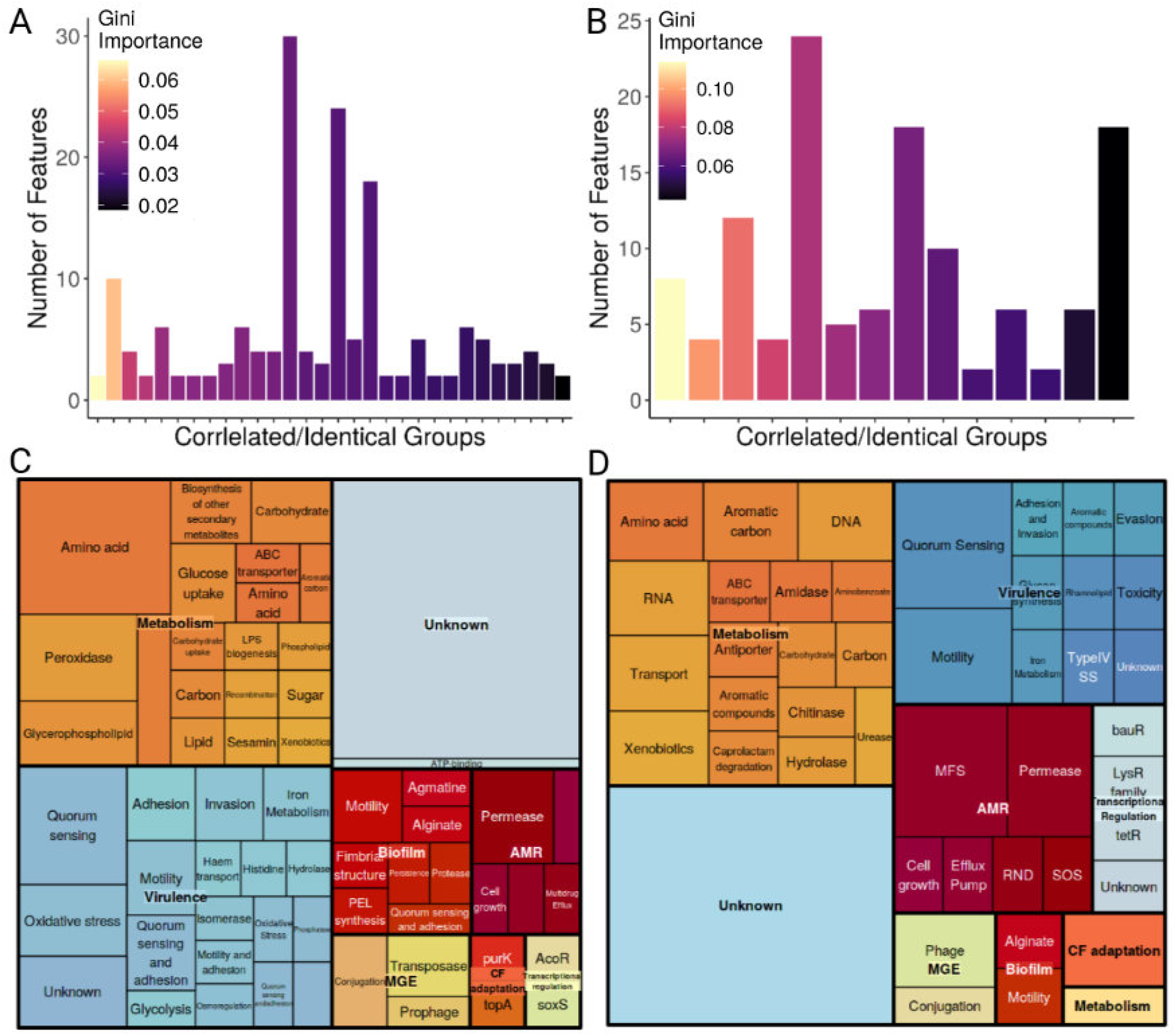
Feature functional annotation. (A) Feature Importance for the PSC control model. The bars indicate the number of features in each of the 30 co-varying group, colored according to the Gini Importance value obtained during random forest training. The features within each group have either identical presence/absence patterns or are highly correlated (Pearson correlation coefficient > 0.7). (B) Same as in A but in this case showing 14 co-varying groups corresponding to the features selected with nPSC. (C) Functional annotation of the PSC selected features including identical and highly correlated patterns. The treemap show the eight functional categories we defined and the subfunctions within. (D) The same for the nPSC selected features.

We found more accessory genes in the PSC selected group, although the difference was not statistically significant. A recombination analysis found that a higher proportion of nPSC unitigs mapped to recombinant genes, but again, the difference was not statistically significant (S7 Fig).

We used annotations from PROKKA [69], MicrobeAnnotator [70], the Virulence Factor Database [71], and the KEGG database [72]. 19% and 15% of mapped genes were identified as virulence factors for the PSC and nPSC pipelines respectively, while 30% and 26% of the genes were annotated as hypothetical proteins.

We further classified the genes into eight functional categories: virulence, biofilm, antimicrobial resistance, CF-associated (genes whose variation has been previously reported to impact Pa adaptation to the CF lung), metabolism, transcriptional regulation, and mobile genetic elements (Fig 9 C and D, Table S2). The top ranked co-variation group for the PSC pipeline was composed of genes of unknown function, while the second group included *pelB*, *bapA*, *ecpE/cup*, and *fliK*, which are genes frequently associated with pathogenicity and CF persistent isolates. PelB is involved in pellicle polysaccharide synthesis, which is an important component of the biofilm structure and mediates tolerance to aminoglycosides [73,74]. BapA is involved in adhesion and biofilm formation [75]. FliK is the flagellar hook length control protein associated with swarming motility abilities [76]. And EcpE/Cup is a fimbrial chaperone in *E. coli* with a role in early stages of biofilm formation and host cell recognition [77].

The third ranked group of covarying features included two non-coding features associated to *aguB*, which encodes a protein responsible for agmatine utilization and associated with biofilm development [78]. Other featured genes found are: virulence-associated *fimA, algI, clpP, nhaP2 popB, hxuA* [79,80]; quorum sensing *qseC, phnB, sasA* [81,82]; iron scavenging *fhuA, pchE* [83–85], as well *purK* and *topA*, which are associated with adaptation to the CF lung [86,87]. One of the selected features that showed a high association to the persistent samples was a non-coding variant between *motY/pyrC*, which is an intergenic region associated with adaptation to the CF lung [88]. Also highly associated to persistent samples were variations in *rhsC, msrB*, which are related to oxidative stress, and *ompU*, which is associated with cell permeability. The nPSC set also contains genes with strong associations to virulence, such as *bamA, adrA* [89,90], *ndvB* and *rpoS* [91], or antibiotic resistance (*umuC, oprM*) [92]. The feature set selected with both models was ranked 16/30 in PSC, 5/14 in nPSC, and showed a strong association with persistent samples. This set of 24 covarying unitigs mapped to *comEC*, *argJ, phbB, pcaK, preA, algI*, and hypothetical proteins, with most mapped to *comEC*, which is a gene necessary for natural transformation in many bacterial taxa [93].

## DISCUSSION

Pa chronic infections in CF pediatric patients have been extensively studied. Pa persistent isolates have been shown to have phenotypic and genotypic characteristics involved in adaptation to the CF lung. Less has been explored on the genetic background of new-onset Pa isolates, usually environmentally acquired, that lead to persistence. In this study, we developed random forest predictive models using a nested cross validation (NCV) design and population structure control (PSC) to predict the outcome of the AET for new-onset Pa infections in children with CF. Our sample consisted of Pa isolates recovered prior to AET, as well as a small set of isolates recovered after AET failure. To handle the vast number of features generated from the genomic variation encoding, we used filters and performed model dependent feature selection.

We were able to reduce the very large number of unitig variants while maintaining predictive power and increasing interpretability, achieving very good predictive performance with only a very small number of unitig features (PSC=30, nPSC=14). Models with and without PSC yield different results, the selected features are different and the prediction error distribution across the phylogeny shows that PSC selected features overcome lineage effects, making them strong candidates to evaluate failure cause.

Individual unitigs showed only weak correlation with the AET outcome, this suggests that multiple variants with small effects play a role in determining AET outcome during early stages of Pa infection. The annotation of high performing unitigs point to different genes with roles in adhesion and motility, as well as biofilm formation, all of these, factors that contribute to reduced effectiveness of antibiotics and are therefore relevant in treatment evasion. The functional redundancy observed in the featured genes imply that there is more than one road that could lead to AET evasion and persistence. We showed that the antibiotic resistance profiles of the isolates recovered from successful and failed AET infections was not significantly different, although post-failure isolates, recovered around a month after AET treatment, do show increased MIC levels for all antibiotics tested except colistin. This increase in antibiotic resistance could be due to reduced permeability or increased expression of efflux pumps as suggested by the functional annotation of unitigs associated to post failure samples.

The use of machine learning to predict traits or outcomes of interest based on genomic data is becoming an increasingly popular and important tool in microbial genomics. While these approaches hold great promise, most studies must find ways to mitigate several intrinsic data characteristics that can negatively impact machine learning model performance. One of these characteristics is the imbalanced sample composition (i.e., many more sample from one outcome or trait class than the other), that if not addressed properly, leads to the under-learning of the minority class. In the worst case, a model can be trained that by chance does not include any representative from the minority class. To overcome this issue, we fixed the class proportions based on their observed values in the total dataset. This ensure that the minority (AET failure) class was adequately represented in both the train and test sets. We also selected the best performing models based on the minority class metrics such as precision and recall.

Another very frequent problem when working with *omics* data is high dimensionality data (i.e., many variables or features) with a small sample size, or the so-called large-p, small-n problem [94]. This problem is particularly challenging when it is difficult or expensive to collect samples, such as is commonly the case in clinical research. This spare dataset problem can lead to ascertainment bias and over-optimistic model performance estimates [95]. We addressed this concern by performing a rarefaction analysis on the unitig to identify the sampling depth needed to interrogate our sequence diversity (S7 Fig). To avoid overfitting, we used NCV and a train/test split approach, this combination produces robust and unbiased estimations [25,95]. Other ways to deal with high dimensionality data include dimensionality reduction and feature selection. We performed dimensionality reduction with MCA and applied filters and wrapping methods on unitig patterns. The former showed good performance but had poor resolution and generates features that are difficult to interpret. Alternatively, our filtering methods and recursive feature elimination approach generated high accuracy models with high interpretability.

An important consideration, particularly when dealing with microbial samples, is population structure, which is non-independence of the samples caused by shared evolutionary history. We addressed this issue by including a population structure control (PSC) that involved blocking the data by BAPS groups or subpopulations when assigning sample to the train or test set. PSC was used exclusively during the NCV along with the feature selection process. This allowed us to measure the impact of blocking and differentiate between features that contribute to, or are responsible for, sample dependence, and those that do not. It is important to make this distinction because we observed that different groups of features can accomplish similar results. Still, we were able to show that the selected features obtained from the PSC modeling were less likely to simply recover the BAPS population structure groups.

Despite the efforts placed into producing models that have high performance and generalization power, they are still based on one-hot encoded sequence diversity, which means that variation found in the population but not in our sample could not be modeled. This is where the mapping and annotation of the selected features (i.e., determining what genes carry the variant feature of interest) becomes of great importance. Even though unseen variation cannot be accounted for, the approach used allows us to identify genes and pathways putatively involved in AET failure and the establishment of chronic infections. It was gratifying to see that many of our PSC selected features mapped to known virulence-associated genes, such as biofilms, iron metabolism and scavenging, motility, and quorum sensing.

In summary, our approach effectively predicted AET failure among new-onset Pa infection in children with CF using Pa genomic sequences. We also showed that including controls for population structure in the analysis is necessary for biological interpretation of features as well as for generalization of failure causes. The power of this approach is made even more evident given the fact that no information on the host factors, comorbidities, or other clinical or environmental data were included in this analysis. These powerful methods provide new avenues for the analysis of high dimensional genomic data and are likely to play a prominent role in predicting complex phenotypes that are underpinned by many polymorphic genes interacting with each other and a suit of environmental and host factors in unpredictable ways.

## METHODS

### Sample Collection

The study cohort consisted of 70 cystic fibrosis pediatric patients from The Hospital for Sick Children, Toronto, Canada, with at least one new-onset Pa infection registered between 2011-2016. Pa isolates were recovered from sputum samples collected before antibiotic treatment and sequenced. If colony morphology variation was observed, different isolates were sequenced to represent the diversity. New-onset Pa infection was defined as first life time acquired infection or a Pa-positive sputum culture at least 12 months. after a previous infection was cleared. AET failure (persistent isolates) was defined as having a positive sputum culture for Pa on the culture done 1 week after completion of AET. Eradicated cases were defined as having a negative sputum culture for Pa in the same time frame. Isolates retrieved from sputum obtained after treatment failure were also sequenced for 10 patients (post-failure samples) to confirm AET outcome assignation. The antibiotic treatment consists of a multi step protocol of inhaled and intravenous tobramycin detailed in [15].

### Antimicrobial susceptibility testing

All Pa isolates were screened for antimicrobial susceptibility by the broth micro-dilution method in accordance with Clinical and Laboratory Standards Institute (CLSI) procedures [96]. Susceptibility profiles were determined for β-lactams (aztreonam, ceftazidime, cefepime, meropenem, imipenem), fluoroquinolones (ciprofloxacin, levofloxacin), aminoglycosides (amikacin, tobramycin, gentamicin), β-lactams/ β-lactamase inhibitor (piperacillin/tazobactam) and colistin. Isolates were grown in Mueller-Hinton II broth overnight at 35°C in a two-fold dilution series of each antibiotic. Results are reported as minimum inhibitory concentration (MIC), and resistant, susceptible or intermediate phenotypes were assigned according to CLSI guidelines.

### Genome analysis

All sequencing was performed on the Illumina NextSeq instrument at the University of Toronto Centre for the Analysis of Genome Evolution and Function (CAGEF). WGS raw data was trimmed using Trimmomatic (Bolger, Lohse, and Usadel 2014)(LEADING:3 TRAILING:3 SLIDINGWINDOW:4:15 MINLEN:80) and assembled with Spades v3.14.1 (--careful -k 21,33,55,77,83,91,101,113,121,127 --mismatch-correction) (Bankevich et al. 2012). Annotation was performed with Prokka (Seemann 2014). Reference genomes were obtained from RefSeq database (PAO1: <u>GCF_000006765.1</u>, PA14: GCF_000404265.1, PA7: GCF_000017205.1) and re-annotated with Prokka. Pangenome analysis were performed using Roary (-i 95 -e --mafft) [97]. RaxML [98] was used for core genome based phylogenetic reconstructions with parameters (-m GTRGAMMA -p 12345 -# 20). Bayesian analysis of population structure were performed using hierBAPS in R with default parameters [99]. Core genome distances were estimated using MASH [100], and accessory genome distances were estimated using bray-curtis dissimilarity index from the gene presence absence matrix obtained with Roary [97].

### Machine learning and model evaluation

#### Encoding genomic variation

We used unitig-counter [62] to create unitigs from the genome assemblies. Unitigs are short sequences of different length that can represent different forms of genetic variation such as SNPs and indels and can dispense with the use of a reference genome. Each genome is represented by a vector of presence/absence of each unitig sequence. A presence-absence matrix for 494 genomes with a total of 542,296 unique unitig patterns was generated. Low and high frequency patterns (<5%, >90%) were removed, a total of 425,005 features remained for further analysis.

#### Feature Extraction - Multiple Corresponding Analysis

Multiple Correspondence Analysis (MCA) was performed on the 425,005 unique unitig patterns matrix using *factominer* library in R [101]. A total of 38 dimensions retaining 80% of the explained variation were kept and used as input features for the model.

#### Feature selection - Model independent selection

Filters were applied on train data only. We calculated Chi-square statistic between each feature and the target and select the desired number of features with best Chi-square scores (p-value < 0.01). Then, we created a correlation matrix with Pearson method and dropped highly correlated features (`Pearson correlation coefficient` > 0.7) leaving 4800 unique unitig patterns for training.

#### Feature selection - Model dependent selection

We used recursive feature elimination (RFE) with a random forest predictor to select features that are more relevant in predicting the target variable. The number of features to select varied between PSC and nPSC pipelines. When using MCA features, we selected 10 features, but when using the filtered unitig one-hot encoded data, we compared independent runs with different numbers of features to select (RFE: n to select= 100,50,30,10).

#### Model design and evaluation

Random forest implementation of scikit-learn (v0.24.1) package in python (v3.6.8) with parameters adjusted during tuning via cross validation was used to build the predictors [102]. We compared two model designs, one consisted of nested cross validation (NCV) with blocking by BAPS groups to control the population structure, and one without blocking. Blocking was applied during the outer loop of the nested cross validation, restricting the use of certain BAPS subpopulations to either the test (15% of the data) or the train (85% of the data) set. We used *GroupShuffleSplit* function to apply blocking taking class proportion into account. Train/test outer split of the models with no PSC were based on class proportion information only (*StratifiedShuffleSplit*).

In every case, during the inner loop the hyperparamter search was performed using the function *GridSearchCV* with cv=3. The hyperparameters combination tested was: n_estimators: 50, 100, 200, class_weight: None, balanced, max_depth: 3, 4, 5, 6, 7, 8. With the best hyperparameter combination chosen, the RFE step is run, where features are selected. The selected features from each outer loop that showed test performance above 0.7 AUC, were combined to fit a new predictor. Performance scores and predictions of each pipeline are obtained at this point.

When using MCA dimensions, the combination of selected features was used to fit a new predictor on whole data using a 10-fold cross-validation. When using one-hot encoded unitig variation, we divided our data set into train and hold-out before the NCV step, therefore selected feature combinations were tested on the held-out dataset and overfitting reduced by further reducing the number of features based on training feature importance.

#### Performance metrics

We used the area under the ROC curve (AUC) for assessing overall performance on training and test sets or training and validation sets. ROC is a probability curve and AUC represents the degree of separability, how much the model is capable of distinguishing between classes. A high AUC indicates low prediction error. Precision, recall and F1-score were estimated for both classes using the *classification_report* function from sickit-learn [102]. Precision measures of how many of the positive predictions made are correct (true positives), recall measures how many positive cases were detected over all the positive cases (sensitivity). F1-score is a single metric that weights precision and recall. The metric used for evaluating feature importance was Gini importance or mean decrease impurity. Permutation importance was also estimated using *permutation_importance* function from scikit-learn (permutations=100).

#### Feature annotation

The one-hot encoded input features represent presence/absence of certain unitig sequences in the genomes. In order to evaluate function and localization of the relevant variation we mapped the unitig sequences to the genomes. The unitigs patterns were mapped using a python script from pyseer [103]. When unitigs mapped onto noncoding regions, the genes up and down stream were identified. All genomes in our database were used as references so that all the patterns could be annotated, and prokka annoatations was improved using MicrobeAnnotator [70], the Virulence Factor Database [71], and the KEGG database [72].

## Data Availability

Genomic sequence data is available from NCBI BioProject ID: PRJNA893599.

## Code Availability

All analysis code is available at https://github.com/luciagrami/EarlyEradication.

## ACKNOWLEDGMENTS

We thank the members of the Guttman lab and the Centre for the Analysis of Genome Evolution and Function (CAGEF) for their assistance and feedback. Special thanks go to Dr. Sylva Donaldson for her project management contributions. The study was supported by a Collaborative Health Research Project grant from the Canadian Institutes of Health Research and the Natural Sciences and Engineering Research Council of Canada (CP-151952).

## SUPPORTING INFORMATION

**S1 Table. Samples used in the study.**

**S2 Table. Features selected with both pipelines.**

**S1 Fig.**
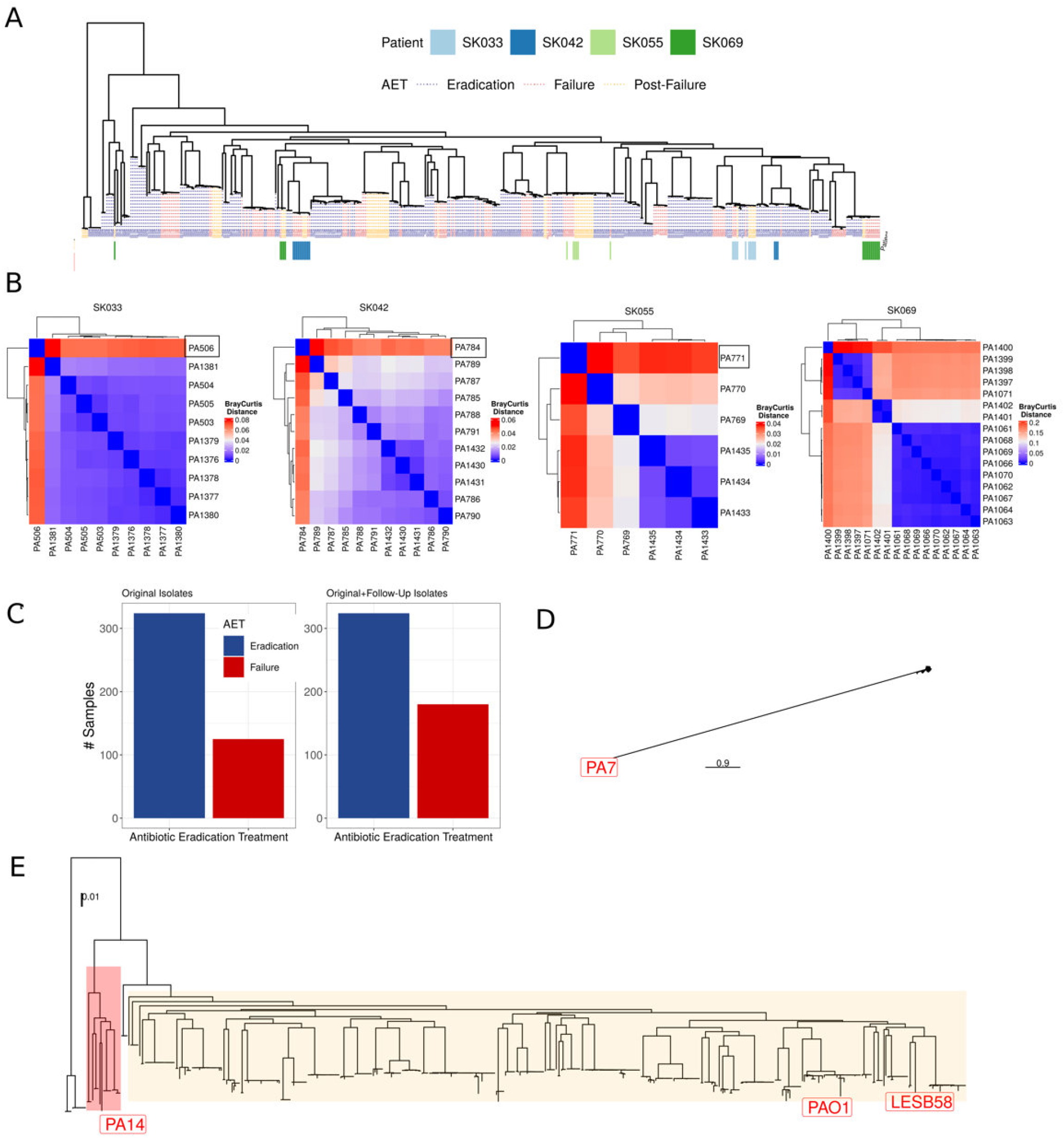
Samples reassignment and phylogenetic diversity. (A) Core genome, midpoint-rooted phylogeny of the 494 Pa strains. AET outcome is represented by the color coding of the dashed lines leading from the terminal nodes to the metadata rows, with blue showing AET success (i.e., eradication), red showing AET failure, and orange showing a post-failure isolate. The annotation bar indicates the patients that showed within infection variation and therefore post-treatment isolates were sequenced. Pre and post treatment samples share a common ancestor, except for one post failure sample, corresponding to patient SK069. (B) Accessory genome distances between pre and post treatment samples. Comparisons were made within patients that showed genetic variation. Bray-Curtis distance was estimated from the gene presence-absence matrix obtained with pangenome analysis. Black squares indicate isolates whose phenotype was modified from failure to eradication due to accessory genome distance. (C) Proportion of samples in each class (eradication and failure) before (left panel) and after (right panel) the samples reassignment and post failure inclusion. (D) Unrooted phylogeny created using the reference genomes PAO1, PA7, LESB58, and PA14. None of our samples were found related to the reference PA7, therefore we removed PA7 to improve the visualization. (E) The phylogeny without PA7 and midpoint rooted. The references, PAO1, PA14 and LESB58 are highlighted.

**S2 Fig.**
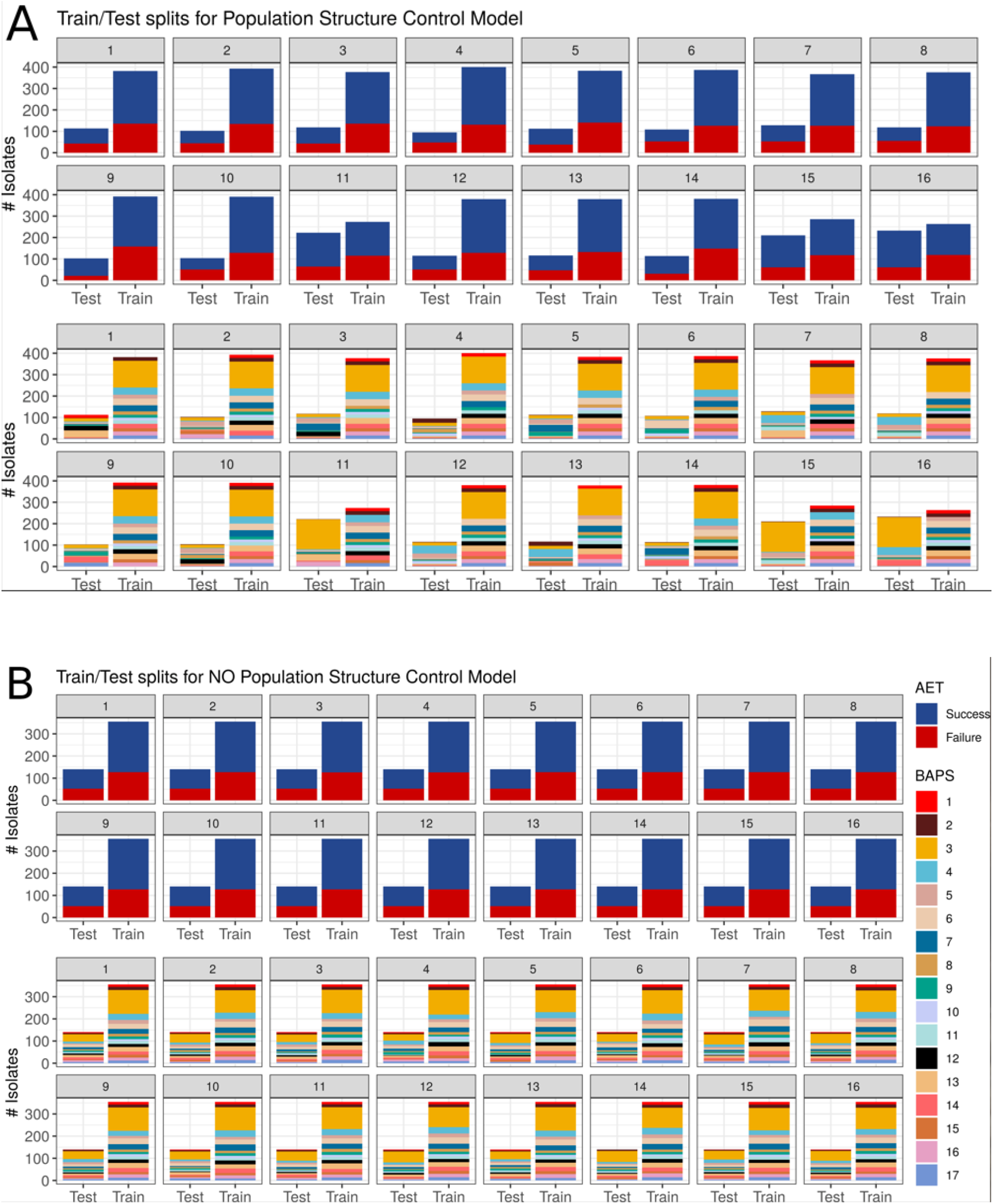
Training/test splits generated using PSC and without PSC during the outer loop of NCV. (A) Train/test splits with population structure control. The top panel displays how treatment outcome (classes) proportions are maintained across all splits. The panel below illustrates the same train/test splits, but this time it can be seen how BAPS subpopulations are distributed. A certain BAPS group can only be in train or test, not in both, conditioning also the size of the splits. (B) Train/test splits without population structure control. The top panel displays how treatment outcome (classes) proportions are maintained across all splits. The panel below shows that without population structure control, a certain BAPS group can be present in both train and test splits.

**S3 Fig.**
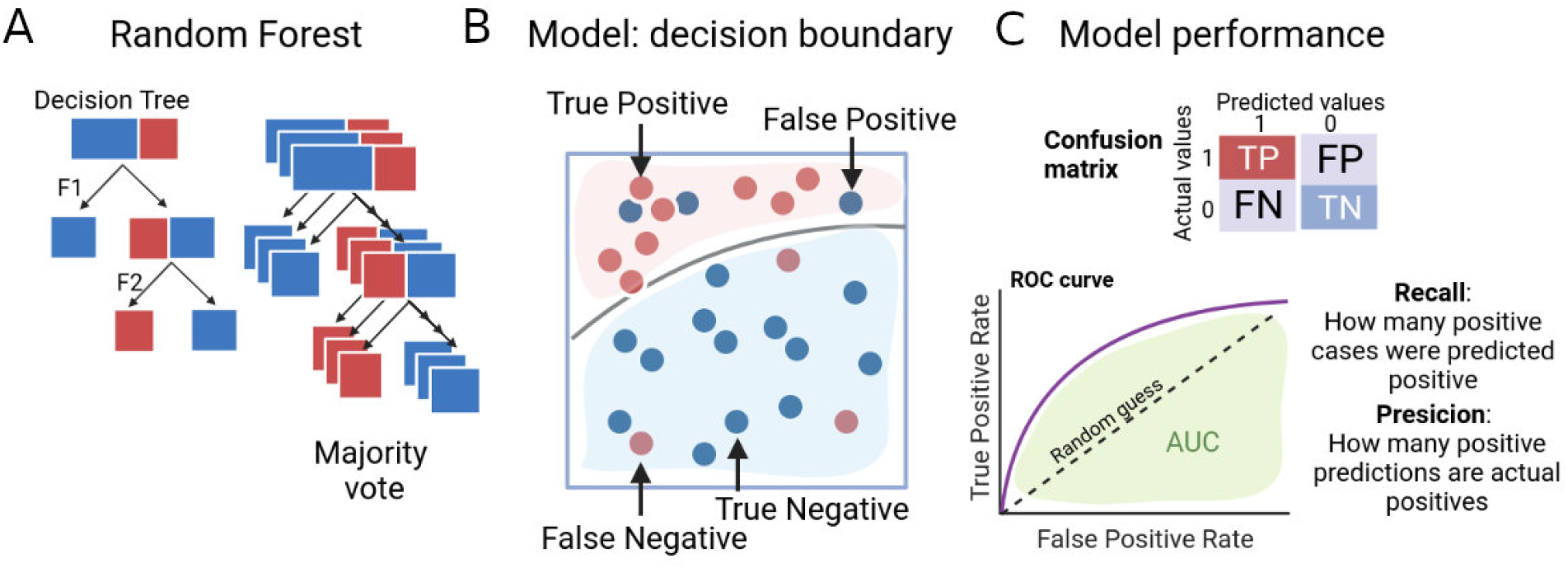
Random Forest design and performance overview. **(A)** Decision trees use tree representations to solve problems, in which leaves represent class labels and internal nodes represent attributes. A random forest is an ensemble of many individual decision trees, each tree’s classification is combined into a final classification through a “majority vote” mechanism. **(B)** A schematic of the decision boundary (partition of the feature space) showing correct and incorrect samples predictions. (C) **Model performance**. In the confusion matrix, the rows represent the true labels and the columns represent the predicted labels. Diagonal values represent the number of times the predicted label matches the true label. Observations in the other cells were mislabeled by the classifier. From the confusion matrix, precision, recall, and F1 score can be derived. A ROC curve (receiver operating characteristic curve) is a graph showing the performance of a classification model at all classification thresholds. This curve plots two parameters, the true and false positive rates. The area under the ROC curve (AUC) provides an aggregate measure of performance across all possible classification thresholds. AUC indicates how well the model can distinguish between classes. Higher the AUC, smaller the prediction error.

**S4 Fig.**
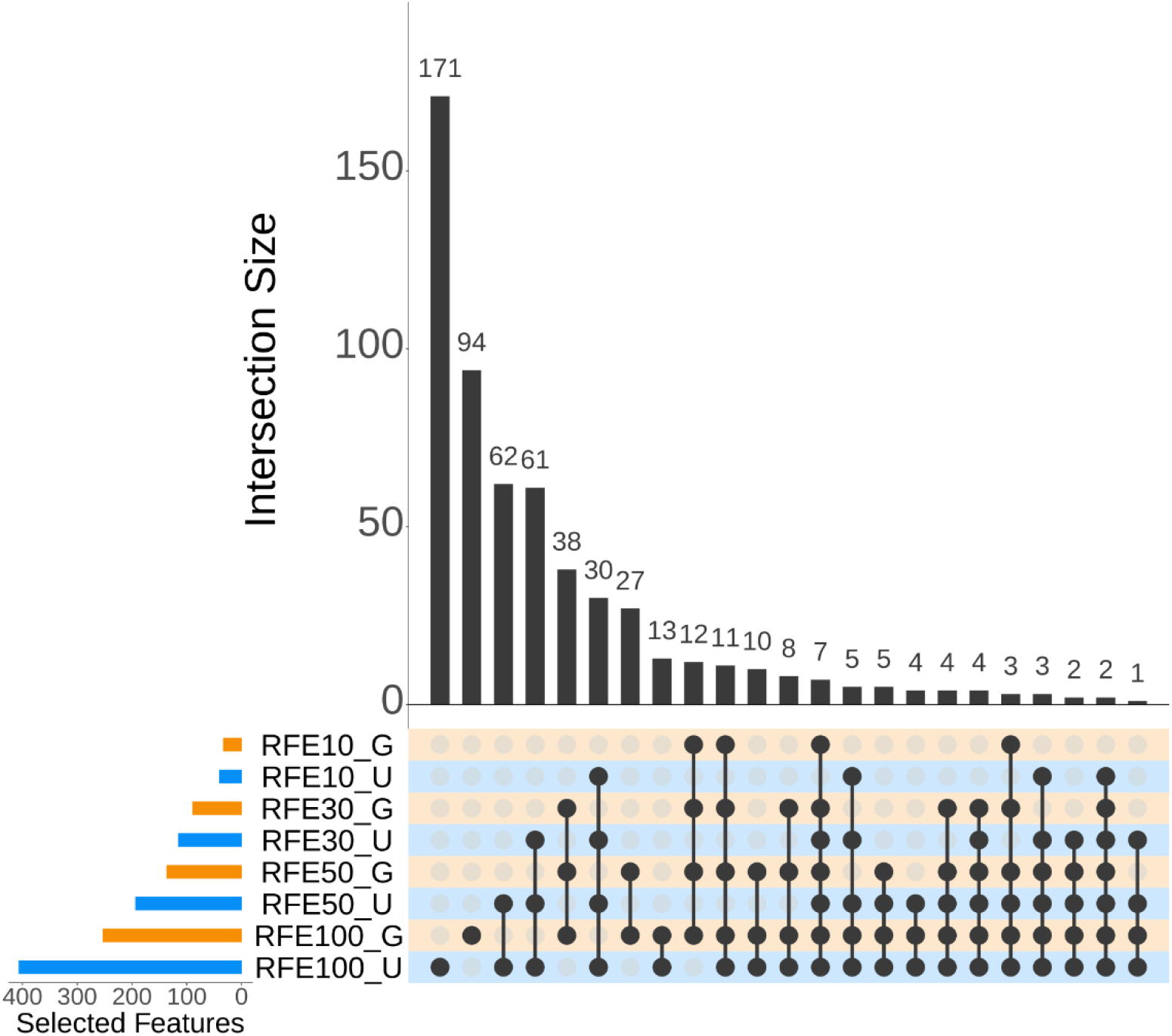
Feature overlap between pipelines with and without population structure control. The intersection plot shows the overlap between features selected during different Recursive Feature Elimination (RFE) instances of the population structure control (PSC) (RFEN_G) and no population structure control (nPSC) (RFEN_U) pipelines. The selected feature set sizes are displayed as horizontal bars on the lower left corner of the image, in blue different RFE instances of the nPSC pipeline, in orange, different RFE instances of the PSC pipeline. Intersection sizes are shown as individual vertical bars. Independent pipelines involved in each intersection are identified with connected black circles under the vertical bars. Unconnected circles represent features that are found exclusively in the corresponding set. Shading in the background of the circles helps differentiate between PSC pipelines (orange) and nPSC pipelines (blue).

**S5 Fig.**
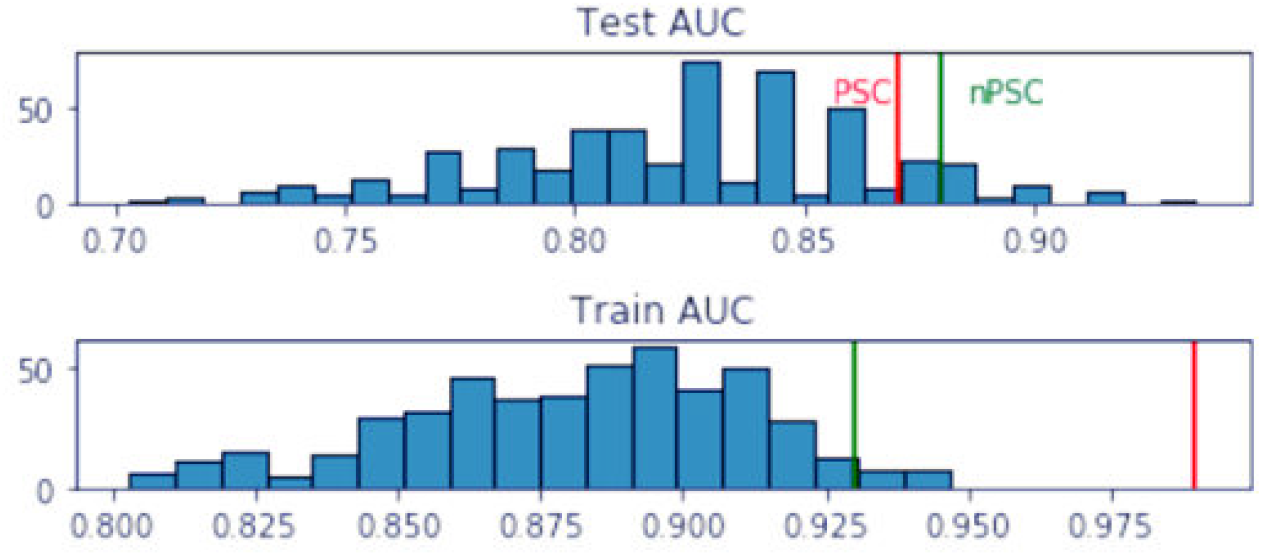
Predictive power of randomly selected features. We fit 500 models with 500 subsets of 25 randomly selected features from the uncorrelated 4800 features set. Test (top panel) and train (bottom) AUC values for the 500 models are shown. In green the AUC values for test and train obtained with the best performing model of the no population structure control pipeline, and in red the test and train values of the best performing model with population structure control. Randomly selected features can show high accuracy most likely due to the correlation of AET outcome and the phylogeny.

**S6 Fig.**
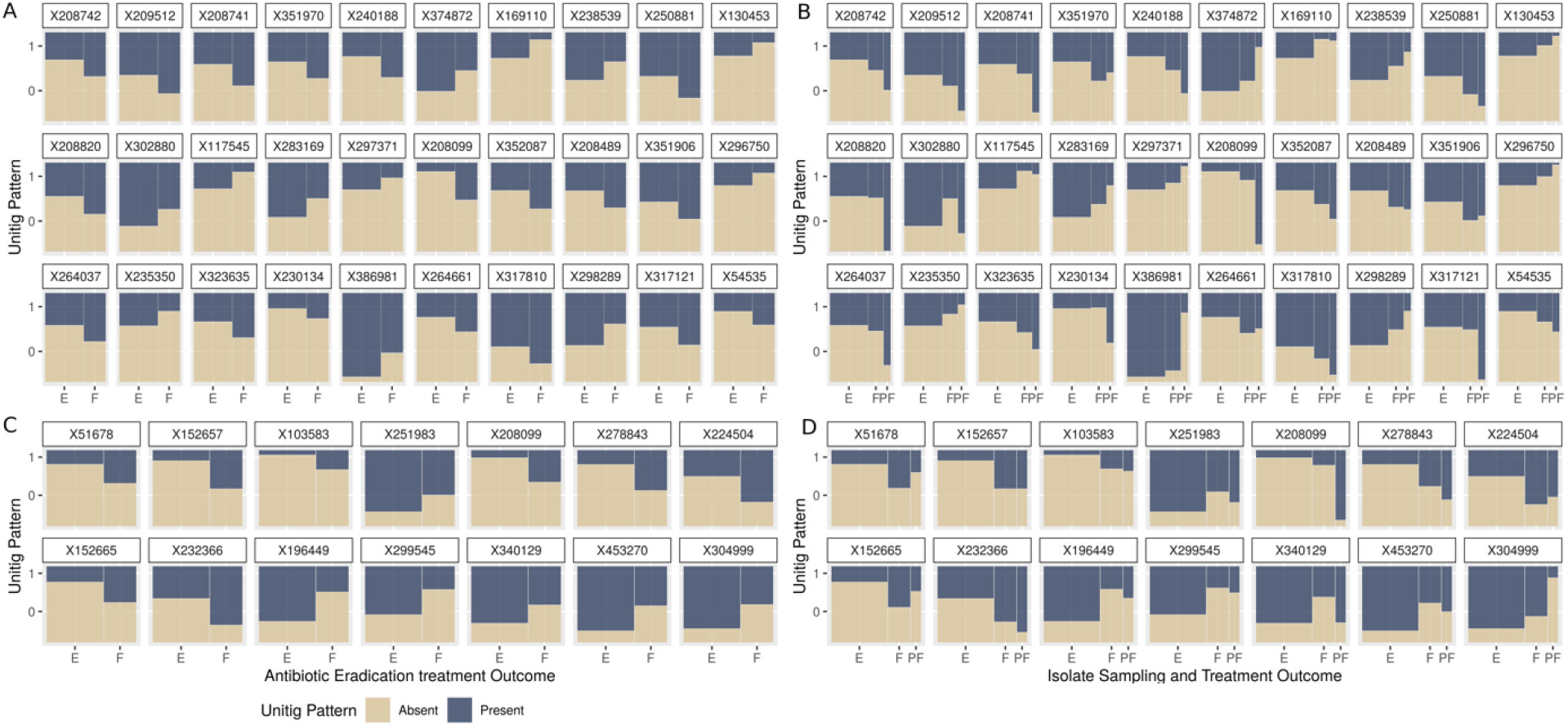
Association of selected features patterns with the Antibiotic Treatment outcome distribution. (A) For each feature selected during the population structure control pipeline (PSC) the comparison of the distribution of the unitig presence/absence pattern in both the eradication and failure groups. (B) The distribution of the unitig presence/absence pattern is now distributed in three groups, eradication, failure and post-failure, to assess the impact of the latter for each independent feature. (C) Same as A for the features selected with the no population structure control pipeline (nPSC). (D) Same as (B) for nPSC selected features.

**S7 Fig.**
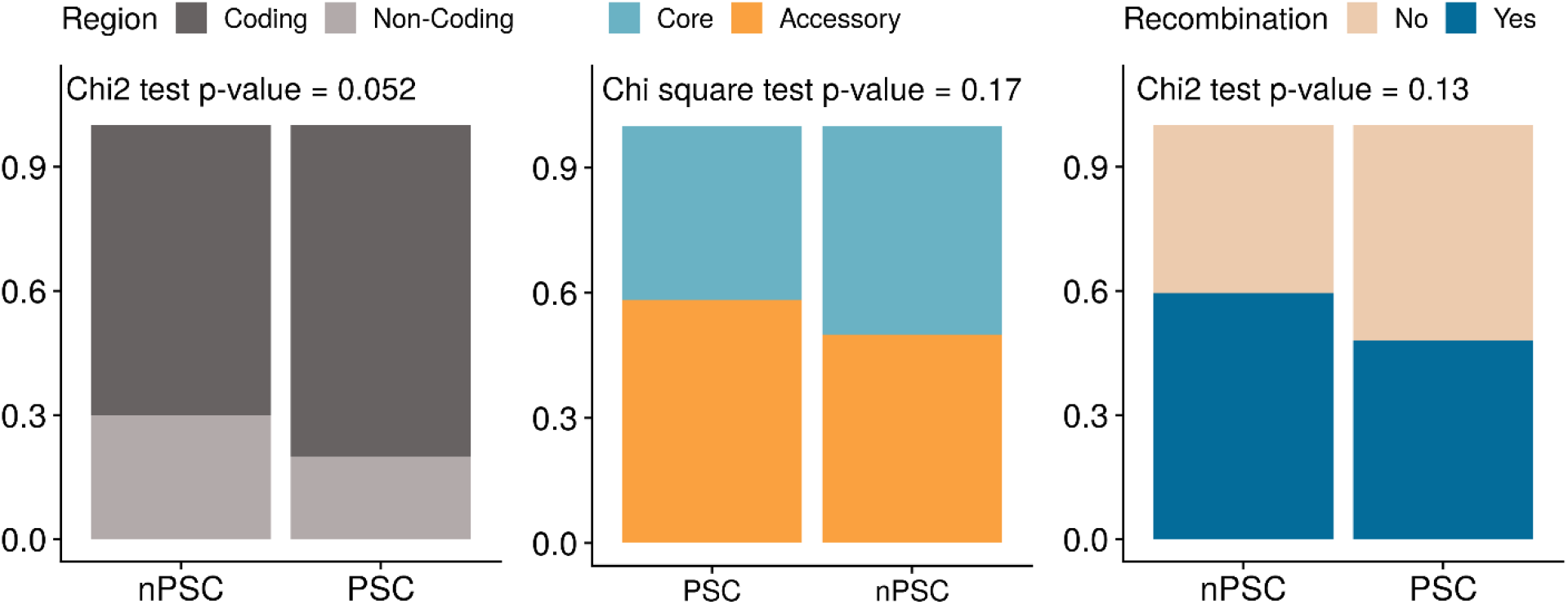
Proportion of features mapping to coding/non-coding regions, accessory and core genes, and genes with/without recombination signal. Chi square association tests were performed in each case.

**S8 Fig.**
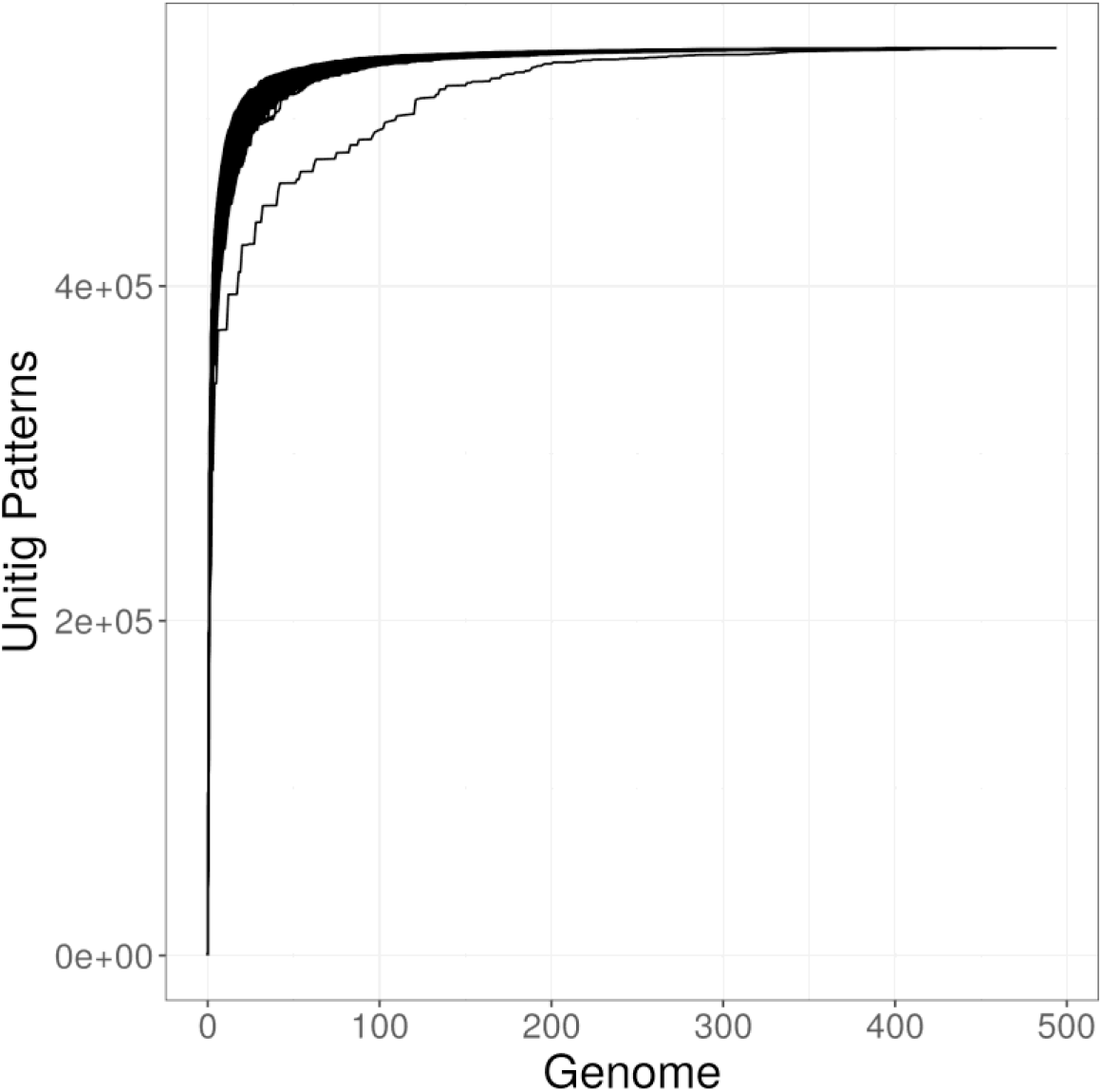
Unitig pattern rarefaction curve.

**S9 Fig.**
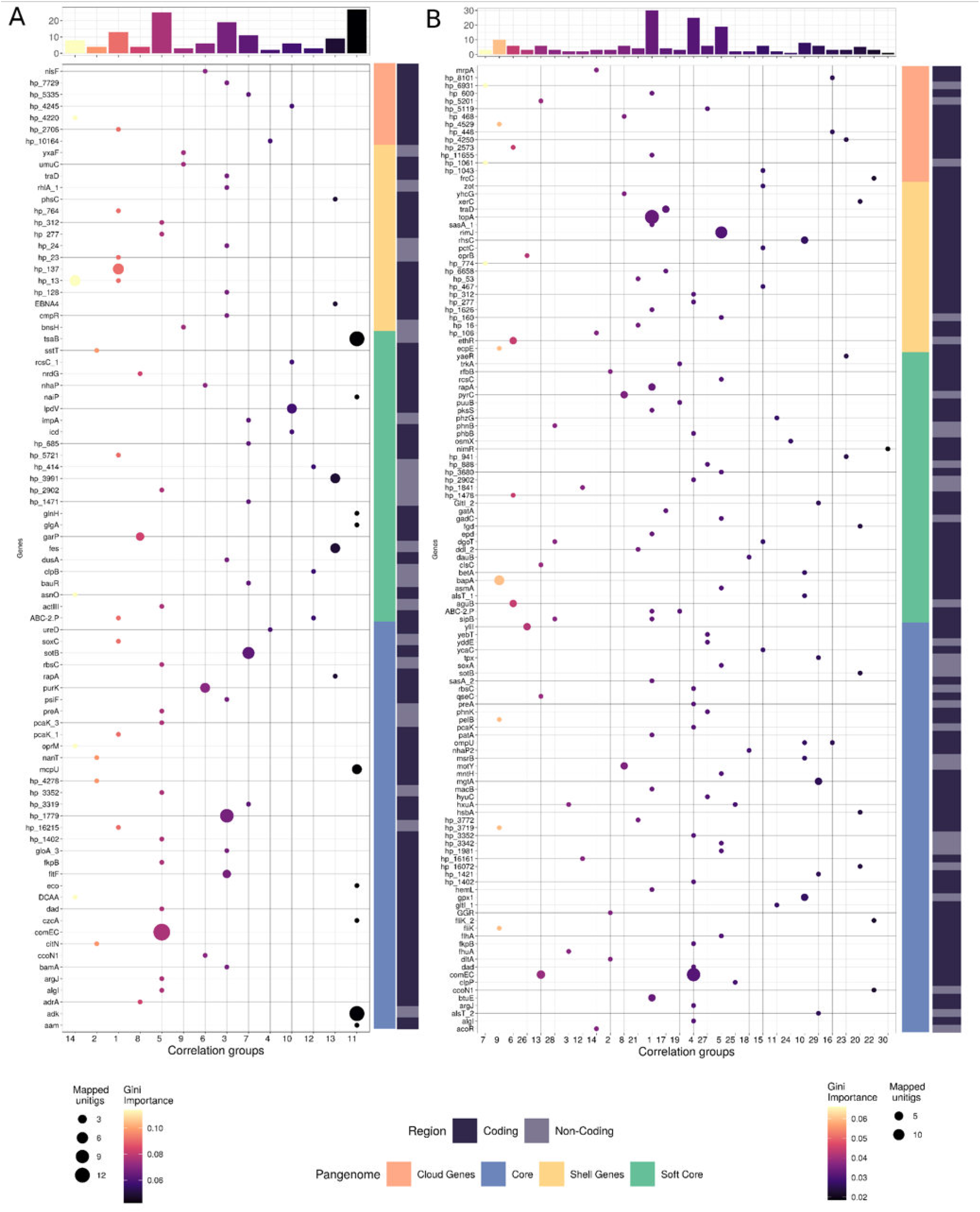
Feature annotation summary. (A) Genes selected with no population structure pipeline (nPSC). The dot plot shows the number of unitigs that mapped to each gene. Gene names are in the y-axis, and covariation groups in the x-axis. Features in one covariation group can map to more than one gene. Features from different covariation groups are less likely to map to the same gene. The top panel shows the Gini importance for the reference feature of the covariation group. Bars and dots are also colored by Gini Importance. The annotations on the right correspond to the pangenome location of the gene, and the nature of the mapped region, coding or non-coding. (B) Genes obtained with the population structure pipeline (PSC). Description is the same as (A).

## REFERENCES

1. Férec C, Scotet V. Genetics of cystic fibrosis: Basics. Arch Pediatr. 2020;27 Suppl 1:eS4–eS7.

2. Bhagirath AY, Li Y, Somayajula D, Dadashi M, Badr S, Duan K. Cystic fibrosis lung environment and *Pseudomonas aeruginosa* infection. BMC Pulm Med. 2016;16(1):174.

3. Rossi E, La Rosa R, Bartell JA, Marvig RL, Haagensen JAJ, Sommer LM, et al. *Pseudomonas aeruginosa* adaptation and evolution in patients with cystic fibrosis. Nat Rev Microbiol. 2021;19(5):331–42.

4. Dickson RP. The microbiome and critical illness. Lancet Respir Med. 2016;4(1):59–72.

5. Whelan FJ, Heirali AA, Rossi L, Rabin HR, Parkins MD, Surette MG. Longitudinal sampling of the lung microbiota in individuals with cystic fibrosis. PLoS One. 2017;12(3):e0172811.

6. Zemanick ET, Wagner BD, Robertson CE, Ahrens RC, Chmiel JF, Clancy JP, et al. Airway microbiota across age and disease spectrum in cystic fibrosis. Eur Respir J. 2017;50(5)

7. Khanolkar RA, Clark ST, Wang PW, Hwang DM, Yau YCW, Waters VJ, et al. Ecological succession of polymicrobial communities in the cystic fibrosis airways. mSystems. 2020;5(6):e00809–20.

8. Coburn B, Wang PW, Diaz Caballero J, Clark ST, Brahma V, Donaldson S, et al. Lung microbiota across age and disease stage in cystic fibrosis. Sci Rep. 2015;5:10241.

9. Davies JC. *Pseudomonas aeruginosa* in cystic fibrosis: pathogenesis and persistence. Paediatr Respir Rev. 2002;3(2):128–34.

10. Scotet V, L’Hostis C, Férec C. The changing epidemiology of cystic fibrosis: incidence, survival and impact of the CFTR gene discovery. Genes (Basel). 2020;11(6)

11. Canada CF. The Canadian cystic fibrosis registry 2020 annual data report. Toronto, Canada: Cystic Fibrosis Canada, 2022.

12. Casaredi IG, Shaw M, Waters V, Seeto R, Blanchard A, Ratjen F. Impact of antibiotic eradication therapy of *Pseudomonas aeruginosa* on long term lung function in cystic fibrosis. J Cyst Fibros. 2022;

13. Mogayzel PJ, Jr., Naureckas ET, Robinson KA, Brady C, Guill M, Lahiri T, et al. Cystic Fibrosis Foundation pulmonary guideline. pharmacologic approaches to prevention and eradication of initial Pseudomonas aeruginosa infection. Ann Am Thorac Soc. 2014;11(10):1640–50.

14. Stanojevic S, Waters V, Mathew JL, Taylor L, Ratjen F. Effectiveness of inhaled tobramycin in eradicating *Pseudomonas aeruginosa* in children with cystic fibrosis. J Cyst Fibros. 2014;13(2):172–8.

15. Blanchard AC, Horton E, Stanojevic S, Taylor L, Waters V, Ratjen F. Effectiveness of a stepwise *Pseudomonas aeruginosa* eradication protocol in children with cystic fibrosis. J Cyst Fibros. 2017;16(3):395–400.

16. Jackson L, Waters V. Factors influencing the acquisition and eradication of early *Pseudomonas aeruginosa* infection in cystic fibrosis. J Cyst Fibros. 2021;20(1):8–16.

17. Morris AJ, Jackson L, Cw Yau Y, Reichhardt C, Beaudoin T, Uwumarenogie S, et al. The role of Psl in the failure to eradicate *Pseudomonas aeruginosa* biofilms in children with cystic fibrosis. NPJ Biofilms Microbiomes. 2021;7(1):63.

18. Moradigaravand D, Palm M, Farewell A, Mustonen V, Warringer J, Parts L. Prediction of antibiotic resistance in *Escherichia coli* from large-scale pan-genome data. PLoS Comput Biol. 2018;14(12):e1006258.

19. Lupolova N, Dallman TJ, Matthews L, Bono JL, Gally DL. Support vector machine applied to predict the zoonotic potential of E. coli O157 cattle isolates. Proc Natl Acad Sci U S A. 2016;113(40):11312–7.

20. Wheeler NE, Gardner PP, Barquist L. Machine learning identifies signatures of host adaptation in the bacterial pathogen *Salmonella enterica*. PLoS Genet. 2018;14(5):e1007333.

21. Khaledi A, Weimann A, Schniederjans M, Asgari E, Kuo TH, Oliver A, et al. Predicting antimicrobial resistance in *Pseudomonas aeruginosa* with machine learning-enabled molecular diagnostics. EMBO Mol Med. 2020;12(3):e10264.

22. Kim JI, Maguire F, Tsang KK, Gouliouris T, Peacock SJ, McAllister TA, et al. Machine learning for antimicrobial resistance prediction: current practice, limitations, and clinical perspective. Clin Microbiol Rev. 2022:e0017921.

23. Nicholls HL, John CR, Watson DS, Munroe PB, Barnes MR, Cabrera CP. Reaching the end-game for GWAS: machine learning approaches for the prioritization of complex disease loci. Front Genet. 2020;11:350.

24. Ritchie MD, Holzinger ER, Li R, Pendergrass SA, Kim D. Methods of integrating data to uncover genotype-phenotype interactions. Nat Rev Genet. 2015;16(2):85–97.

25. Whalen S, Schreiber J, Noble WS, Pollard KS. Navigating the pitfalls of applying machine learning in genomics. Nat Rev Genet. 2022;23(3):169–81.

26. Hicks AL, Wheeler N, Sanchez-Buso L, Rakeman JL, Harris SR, Grad YH. Evaluation of parameters affecting performance and reliability of machine learning-based antibiotic susceptibility testing from whole genome sequencing data. PLoS Comput Biol. 2019;15(9):e1007349.

27. de Los Campos G, Vazquez AI, Hsu S, Lello L. Complex-trait prediction in the era of big data. Trends Genet. 2018;34(10):746–54.

28. Womack JE, Jang HJ, Lee MO. Genomics of complex traits. Ann N Y Acad Sci. 2012;1271(1):33–6.

29. Glazier AM, Nadeau JH, Aitman TJ. Finding genes that underlie complex traits. Science. 2002;298(5602):2345–9.

30. Hirschhorn JN, Daly MJ. Genome-wide association studies for common diseases and complex traits. Nat Rev Genet. 2005;6(2):95–108.

31. Allen JP, Snitkin E, Pincus NB, Hauser AR. Forest and trees: exploring bacterial virulence with genome-wide association studies and machine learning. Trends Microbiol. 2021;

32. Falush D. Bacterial genomics: Microbial GWAS coming of age. Nat Microbiol. 2016;1:16059.

33. Falush D, Bowden R. Genome-wide association mapping in bacteria? Trends Microbiol. 2006;14(8):353–5.

34. Power RA, Parkhill J, de Oliveira T. Microbial genome-wide association studies: lessons from human GWAS. Nat Rev Genet. 2017;18(1):41–50.

35. Hellwege JN, Keaton JM, Giri A, Gao X, Velez Edwards DR, Edwards TL. Population stratification in genetic association studies. Curr Protoc Hum Genet. 2017;95:1.22.1–1..3.

36. Vilhjálmsson BJ, Nordborg M. The nature of confounding in genome-wide association studies. Nat Rev Genet. 2013;14(1):1–2.

37. Camacho DM, Collins KM, Powers RK, Costello JC, Collins JJ. Next-generation machine learning for biological networks. Cell. 2018;173(7):1581–92.

38. Nguyen TT, Huang JZ, Nguyen TT. Unbiased feature selection in learning random forests for high-dimensional data. ScientificWorldJournal. 2015;2015:471371.

39. Chen X, Ishwaran H. Random forests for genomic data analysis. Genomics. 2012;99(6):323–9.

40. Montesinos López OA, Montesinos López A, Crossa J. Random Forest for Genomic Prediction. Multivariate statistical machine learning methods for genomic prediction. Cham: Springer International Publishing; 2022. p. 633–81.

41. Saarela M, Jauhiainen S. Comparison of feature importance measures as explanations for classification models. SN Applied Sciences. 2021;3(2):272.

42. Parvandeh S, Yeh HW, Paulus MP, McKinney BA. Consensus features nested cross-validation. Bioinformatics. 2020;36(10):3093–8.

43. Tadist K, Najah S, Nikolov NS, Mrabti F, Zahi A. Feature selection methods and genomic big data: a systematic review. Journal of Big Data. 2019;6(1):79.

44. Vidya P, Smith L, Beaudoin T, Yau YC, Clark S, Coburn B, et al. Chronic infection phenotypes of *Pseudomonas aeruginosa* are associated with failure of eradication in children with cystic fibrosis. Eur J Clin Microbiol Infect Dis. 2016;35(1):67–74.

45. Lyczak JB, Cannon CL, Pier GB. Lung infections associated with cystic fibrosis. Clin Microbiol Rev. 2002;15(2):194–222.

46. Muhlebach MS, Zorn BT, Esther CR, Hatch JE, Murray CP, Turkovic L, et al. Initial acquisition and succession of the cystic fibrosis lung microbiome is associated with disease progression in infants and preschool children. PLoS Pathog. 2018;14(1):e1006798.

47. Brown PS, Pope CE, Marsh RL, Qin X, McNamara S, Gibson R, et al. Directly sampling the lung of a young child with cystic fibrosis reveals diverse microbiota. Ann Am Thorac Soc. 2014;11(7):1049–55.

48. Frayman KB, Armstrong DS, Carzino R, Ferkol TW, Grimwood K, Storch GA, et al. The lower airway microbiota in early cystic fibrosis lung disease: a longitudinal analysis. Thorax. 2017;72(12):1104–12.

49. Ozer EA, Nnah E, Didelot X, Whitaker RJ, Hauser AR. The population structure of *Pseudomonas aeruginosa* is characterized by genetic isolation of exoU+ and exoS+ lineages. Genome Biol Evol. 2019;11(1):1780–96.

50. Lees JA, Mai TT, Galardini M, Wheeler NE, Horsfield ST, Parkhill J, et al. Improved prediction of bacterial genotype-phenotype associations using interpretable pangenome-spanning regressions. mBio. 2020;11(4)

51. Chen PE, Shapiro BJ. The advent of genome-wide association studies for bacteria. Curr Opin Microbiol. 2015;25:17–24.

52. Dutilh BE, Backus L, Edwards RA, Wels M, Bayjanov JR, van Hijum SA. Explaining microbial phenotypes on a genomic scale: GWAS for microbes. Brief Funct Genomics. 2013;12(4):366–80.

53. Cheng L, Connor TR, Sirén J, Aanensen DM, Corander J. Hierarchical and spatially explicit clustering of DNA sequences with BAPS software. Mol Biol Evol. 2013;30(5):1224–8.

54. Corander J, Marttinen P, Siren J, Tang J. Enhanced Bayesian modelling in BAPS software for learning genetic structures of populations. BMC Bioinformatics. 2008;9:539.

55. Tang J, Hanage WP, Fraser C, Corander J. Identifying currents in the gene pool for bacterial populations using an integrative approach. PLoS Comput Biol. 2009;5(8):e1000455.

56. Tonkin-Hill G, Lees JA, Bentley SD, Frost SDW, Corander J. Fast hierarchical Bayesian analysis of population structure. Nucleic Acids Res. 2019;47(11):5539–49.

57. Armstrong D, Bell S, Robinson M, Bye P, Rose B, Harbour C, et al. Evidence for spread of a clonal strain of *Pseudomonas aeruginosa* among cystic fibrosis clinics. J Clin Microbiol. 2003;41(5):2266–7.

58. Spencker FB, Haupt S, Claros MC, Walter S, Lietz T, Schille R, et al. Epidemiologic characterization of *Pseudomonas aeruginosa* in patients with cystic fibrosis. Clin Microbiol Infect. 2000;6(11):600–7.

59. Benkwitz-Bedford S, Palm M, Demirtas TY, Mustonen V, Farewell A, Warringer J, et al. Machine learning prediction of resistance to subinhibitory antimicrobial concentrations from *Escherichia coli* genomes. mSystems. 2021;6(4):e0034621.

60. Pesesky MW, Hussain T, Wallace M, Patel S, Andleeb S, Burnham CD, et al. Evaluation of machine learning and rules-based approaches for predicting antimicrobial resistance profiles in gram-negative bacilli from whole genome sequence data. Front Microbiol. 2016;7:1887.

61. Stoesser N, Batty EM, Eyre DW, Morgan M, Wyllie DH, Del Ojo Elias C, et al. Predicting antimicrobial susceptibilities for *Escherichia coli* and *Klebsiella pneumoniae* isolates using whole genomic sequence data. J Antimicrob Chemother. 2013;68(10):2234–44.

62. Jaillard M, Lima L, Tournoud M, Mahe P, van Belkum A, Lacroix V, et al. A fast and agnostic method for bacterial genome-wide association studies: Bridging the gap between k-mers and genetic events. PLoS Genet. 2018;14(11):e1007758.

63. Arning N, Sheppard SK, Bayliss S, Clifton DA, Wilson DJ. Machine learning to predict the source of campylobacteriosis using whole genome data. PLoS Genet. 2021;17(10):e1009436.

64. Sarica A, Cerasa A, Quattrone A. Random forest algorithm for the classification of neuroimaging data in Alzheimer’s disease: a systematic review. Front Aging Neurosci. 2017;9:329.

65. Breiman L. Random forests. Machine Learning. 2001;45(1):5–32.

66. Biau G. Analysis of a random forests model. J Mach Learn Res. 2012;13(null):1063–95.

67. Guyon I, Elisseeff A. An introduction to variable and feature selection. J Mach Learn Res. 2003;3(7-8):1157–82.

68. Le Roux B, Rouanet H. Geometric data analysis: from correspondence analysis to structured data analysis: Springer Science & Business Media; 2004.

69. Seemann T. Prokka: rapid prokaryotic genome annotation. Bioinformatics. 2014;30(14):2068–9.

70. Ruiz-Perez CA, Conrad RE, Konstantinidis KT. MicrobeAnnotator: a user-friendly, comprehensive functional annotation pipeline for microbial genomes. BMC Bioinformatics. 2021;22(1):11.

71. Chen L, Yang J, Yu J, Yao Z, Sun L, Shen Y, et al. VFDB: a reference database for bacterial virulence factors. Nucleic Acids Res. 2005;33(Database issue):D325–8.

72. Kanehisa M, Goto S. KEGG: kyoto encyclopedia of genes and genomes. Nucleic Acids Res. 2000;28(1):27–30.

73. Marmont LS, Whitfield GB, Rich JD, Yip P, Giesbrecht LB, Stremick CA, et al. PelA and PelB proteins form a modification and secretion complex essential for Pel polysaccharide-dependent biofilm formation in *Pseudomonas aeruginosa*. J Biol Chem. 2017;292(47):19411–22.

74. Friedman L, Kolter R. Two genetic loci produce distinct carbohydrate-rich structural components of the *Pseudomonas aeruginosa* biofilm matrix. J Bacteriol. 2004;186(14):4457–65.

75. de Bentzmann S, Giraud C, Bernard CS, Calderon V, Ewald F, Plésiat P, et al. Unique biofilm signature, drug susceptibility and decreased virulence in *Drosophila* through the *Pseudomonas aeruginosa* two-component system PprAB. PLoS Pathog. 2012;8(11):e1003052.

76. Waters RC, O’Toole PW, Ryan KA. The FliK protein and flagellar hook-length control. Protein Sci. 2007;16(5):769–80.

77. Berne C, Ducret A, Hardy GG, Brun YV. Adhesins Involved in Attachment to Abiotic Surfaces by Gram-Negative Bacteria. Microbiol Spectr. 2015;3(4)

78. Williams BJ, Du RH, Calcutt MW, Abdolrasulnia R, Christman BW, Blackwell TS. Discovery of an operon that participates in agmatine metabolism and regulates biofilm formation in *Pseudomonas aeruginosa*. Mol Microbiol. 2010;76(1):104–19.

79. Horna G, Ruiz J. Type 3 secretion system of *Pseudomonas aeruginosa*. Microbiol Res. 2021;246:126719.

80. Otero-Asman JR, García-García AI, Civantos C, Quesada JM, Llamas MA. *Pseudomonas aeruginosa* possesses three distinct systems for sensing and using the host molecule haem. Environ Microbiol. 2019;21(12):4629–47.

81. Jones CJ, Grotewold N, Wozniak DJ, Gloag ES. *Pseudomonas aeruginosa* initiates a rapid and specific transcriptional response during surface attachment. J Bacteriol. 2022;204(5):e0008622.

82. Wang C, Chen W, Xia A, Zhang R, Huang Y, Yang S, et al. Carbon starvation induces the expression of PprB-regulated genes in *Pseudomonas aeruginosa*. Appl Environ Microbiol. 2019;85(22)

83. Cunrath O, Gasser V, Hoegy F, Reimmann C, Guillon L, Schalk IJ. A cell biological view of the siderophore pyochelin iron uptake pathway in *Pseudomonas aeruginosa*. Environ Microbiol. 2015;17(1):171–85.

84. Gi M, Lee KM, Kim SC, Yoon JH, Yoon SS, Choi JY. A novel siderophore system is essential for the growth of *Pseudomonas aeruginosa* in airway mucus. Sci Rep. 2015;5:14644.

85. Moynié L, Luscher A, Rolo D, Pletzer D, Tortajada A, Weingart H, et al. Structure and Function of the PiuA and PirA Siderophore-Drug Receptors from *Pseudomonas aeruginosa* and *Acinetobacter baumannii*. Antimicrob Agents Chemother. 2017;61(4)

86. Kumar SS, Penesyan A, Elbourne LDH, Gillings MR, Paulsen IT. Catabolism of nucleic acids by a cystic fibrosis *Pseudomonas aeruginosa* isolate: an adaptive pathway to cystic fibrosis sputum environment. Front Microbiol. 2019;10:1199.

87. La Rosa R, Rossi E, Feist AM, Johansen HK, Molin S. Compensatory evolution of *Pseudomonas aeruginosa’s* slow growth phenotype suggests mechanisms of adaptation in cystic fibrosis. Nat Commun. 2021;12(1):3186.

88. Khademi SMH, Sazinas P, Jelsbak L. Within-host adaptation mediated by intergenic evolution in *Pseudomonas aeruginosa*. Genome Biol Evol. 2019;11(5):1385–97.

89. Hoang HH, Nickerson NN, Lee VT, Kazimirova A, Chami M, Pugsley AP, et al. Outer membrane targeting of *Pseudomonas aeruginosa* proteins shows variable dependence on the components of Bam and Lol machineries. mBio. 2011;2(6)

90. Muriel C, Blanco-Romero E, Trampari E, Arrebola E, Durán D, Redondo-Nieto M, et al. The diguanylate cyclase AdrA regulates flagellar biosynthesis in *Pseudomonas fluorescens* F113 through SadB. Sci Rep. 2019;9(1):8096.

91. Hall CW, Hinz AJ, Gagnon LB, Zhang L, Nadeau JP, Copeland S, et al. *Pseudomonas aeruginosa* Biofilm Antibiotic Resistance Gene ndvB Expression Requires the RpoS Stationary-Phase Sigma Factor. Appl Environ Microbiol. 2018;84(7)

92. Migliorini LB, Brüggemann H, de Sales RO, Koga PCM, de Souza AV, Martino MDV, et al. Mutagenesis induced by sub-lethal doses of ciprofloxacin: genotypic and phenotypic differences between the *Pseudomonas aeruginosa* strain PA14 and clinical isolates. Front Microbiol. 2019;10:1553.

93. Pimentel ZT, Zhang Y. Evolution of the natural transformation protein, ComEC, in bacteria. Front Microbiol. 2018;9:2980.

94. Huynh P-H, Nguyen VH, Do T-N. Improvements in the Large p, Small n Classification Issue. SN Computer Science. 2020;1(4):207.

95. Vabalas A, Gowen E, Poliakoff E, Casson AJ. Machine learning algorithm validation with a limited sample size. PLoS One. 2019;14(11):e0224365.

96. CLSI. Methods for dilution antimicrobial susceptibility tests for bacteria that grow aerobically; approved standard - ninth edition. CLSI document 2012:M07–A9.

97. Page AJ, Cummins CA, Hunt M, Wong VK, Reuter S, Holden MT, et al. Roary: rapid large-scale prokaryote pan genome analysis. Bioinformatics. 2015;31(22):3691–3.

98. Stamatakis A. RAxML-VI-HPC: maximum likelihood-based phylogenetic analyses with thousands of taxa and mixed models. Bioinformatics. 2006;22(21):2688–90.

99. Tonkin-Hill G, Lees JA, Bentley SD, Frost SDW, Corander J. RhierBAPS: An R implementation of the population clustering algorithm hierBAPS. Wellcome Open Res. 2018;3:93.

100. Ondov BD, Treangen TJ, Melsted P, Mallonee AB, Bergman NH, Koren S, et al. Mash: fast genome and metagenome distance estimation using MinHash. Genome Biol. 2016;17(1):132.

101. Lê S, Josse J, Husson F. FactoMineR: an R package for multivariate analysis. Journal of Statistical Software. 2008;25:1–18.

102. Hao J, Ho TK. Machine learning made easy: a review of scikit-learn package in python programming language. Journal of Educational and Behavioral Statistics. 2019;44(3):348–61.

103. Lees JA, Galardini M, Bentley SD, Weiser JN, Corander J. pyseer: a comprehensive tool for microbial pangenome-wide association studies. Bioinformatics. 2018;34(24):4310–2.

